# Hundreds of independent midsize deletions mediate DNA loss in wild relatives of Red Jungle Fowl

**DOI:** 10.1101/2023.07.18.549606

**Authors:** Ashutosh Sharma, Sagar Sharad Shinde, Nagarjun Vijay

## Abstract

Small and midsize deletions and insertions (InDels) are major events that play a crucial role in the evolution of genome size and contribute to the genetic and phenotypic diversity of species. In recent years, considerable attention has been given to studying small indels associated with various developmental, growth, and production traits in domestic chicken breeds. Additionally, small and midsize indels have been studied between chicken and phylogenetically more distant species such as duck, turkey, rock pigeon, and other passerine birds. However, the investigation of small and midsize deletions in the wild relatives of chickens has been relatively overlooked until now. To address this gap, our study aimed to identify the presence and distribution of midsize deletions (> 1 Kb) in the wild relatives of chickens. We conducted a comparative genomic analysis using high-quality genomic data from four species belonging to the *Gallus* genus. Our analysis revealed the existence of more than 125 midsize deletions in the three other species compared to *Gallus gallus* (red junglefowl). These midsize deletions were found to be distributed in intergenic regions and within introns of various protein-coding genes but not in the exonic regions of protein-coding genes. Furthermore, we observed a trend between the number of midsize deletions and the phylogenetic distance in the phylogeny of the *Gallus* genus. The most ancestral species, *Gallus varius* (green junglefowl), exhibited the highest deletions, followed by *Gallus lafayettii* (Ceylon junglefowl) and *Gallus sonneratii* (grey junglefowl). Some protein-coding genes harboring deletions in their introns and upstream regions were associated with body development, production, growth traits, abdominal fat deposition, behavioral patterns such as stress, fear, anxiety, plumage color, and adaptation to extreme climatic conditions. Our study finds that the midsize deletions identified in wild relatives of red junglefowl contribute less than 1% of DNA loss with a rate of 8-44 Kb/My during the evolution of the *Gallus* genus.

## Introduction

The dynamics of DNA loss and gain through various independent processes determines the trajectory of genome size evolution. Insertion of transposable elements (TE), repeat expansion/contraction, polyploidization, and insertion/deletion events of different sizes are prominent mechanisms of DNA loss and gain (Ren et al., 2018; Thomas et al., 2003). Genome size expansion through repeat accumulation has been studied in diverse species (Blommaert, 2020; Lehmann et al., 2021; Wang et al., 2019; Wu et al., 2000). Empirical studies have documented that TE sequences are a substantial factor in the genome size variation in plants and animals (Elliott and Gregory, 2015; Lee and Kim, 2014; Zuccolo et al., 2007). The percentage of TE content positively correlates with genome size, where larger genomes have a greater fraction of TE and vice versa for smaller genomes. In addition to TE, the rate of DNA loss through small deletions and insertions can influence genome size variation and is a key parameter for studying genome size evolution (Petrov, 2002a). The Mutational Equilibrium Model of genome size evolution suggests that the ratio of DNA loss to DNA gain determines the genome size equilibrium. Large segmental deletions can cause deleterious effects compared to small insertions and deletions. In contrast, large insertions also have a powerful impact, not necessarily deleterious, increasing genome size. Hence, the mutational equilibrium model proposes that DNA loss predominantly occurs through small deletions, while DNA gains primarily result from large insertions (Petrov, 2002b). The genomic rate of both processes increases at a different degree with an increase in genome size. DNA loss occurs linearly in smaller genomes, and DNA gain outpaces DNA loss, increasing genome size. However, in the case of larger genomes, the DNA loss is more rapid than the gain resulting in genome size equilibrium.

The more recent “accordion” model suggests genome size evolution occurs through DNA loss via large segmental deletions, which is counteracted by DNA gain through TE expansion. In the context of the evolution of avian and eutherian taxa, it has been observed that small or microdeletions (1-30 bp) make a minimal contribution to DNA loss and genome size evolution. Conversely, midsize deletions (30 bp - 10 Kb) play a more pivotal role in DNA loss. On average, micro and midsize deletions account for approximately 20% of DNA loss, indicating that large segmental deletions (>10 Kb) are the primary events responsible for the majority of DNA loss in avian and eutherian taxa (Kapusta et al., 2017). Large segmental deletions of size 1511 Kb, 845 Kb (Nóbrega et al., 2004), and 31 Kb have been reported in mammals in previous studies (Han et al., 2007; McLean and Bejerano, 2008). In birds, 118 regions collectively spanning 58 Mb are deleted through large segmental deletions compared to reptiles (the largest is 2.1 Mb compared to green anole). These large segmental deletions have resulted in the loss of protein-coding genes in birds and may have contributed to their compact genome size and specific traits (Zhang et al., 2014). Microdeletions can determine gene structure, expression, pre-mRNA splicing, and chromosomal rearrangement (Fontanillas et al., 2007) with advantageous or deleterious effects on the genome leading to alterations in phenotypes. The impact of deletions on gene function depends on its location within the genome, such as intergenic regions, exons, introns, and regulatory regions (Yan et al., 2014). Deletions in coding regions can result in frameshift mutations, leading to gene truncation or neofunctionalization. Likewise, deletions in regulatory or intergenic regions can disrupt motifs or promoter-binding sites (Zhou et al., 2010). Reduction in genome size through deletions occurs at a specific size and region (intergenic and non-coding) until it affects the fitness of species (Petrov, 2002b).

Birds and mammals, the two major taxa, exhibit significant interspecific variations in genome size. Birds have a genome size ranging from approximately 1 to 2.1 Gb, while mammals have a wider range of genome sizes from around 1.6 to 6.3 Gb (Kapusta et al., 2017). The avian genome size is highly constrained, with an average size of 1.4 pg. and only a 2-fold variation in the entire class (Gregory T., 2005). Compared to other tetrapod classes, avian genomes have a smaller mean size and lower variation. The conserved and reduced genome size observed in birds suggests a potential monophyletic origin or evolutionary constraint related to the metabolic demands of flight (Tiersch and Wachtel, 1991). Substantial evidence supports a correlation between flight evolution and genome size reduction (Boschiero et al., 2015). Conversely, the loss or reduced flight ability in certain bird species is associated with increased genome size. The decrease in genome size has been influenced by ongoing purifying selection, which becomes relaxed in flightless or ratite birds (Hughes and Friedman, 2008). From an evolutionary perspective, the reduction in genome size compared to an ancestor with a large genome could be attributed to a higher rate of loss of single-copy sequences compared to repetitive sequences (Tiersch and Wachtel, 1991). However, our understanding of the mechanisms underlying the loss of single-copy DNA sequences through micro, midsize, and large segmental deletions is limited due to the technical challenges associated with accurately identifying deletion events. The distinct roles of deletions of different sizes still require further elucidation, which can be achieved by using high-quality genome assemblies and short and long-read genomic data.

Red junglefowl (*Gallus gallus*) is one of the most important domestic animals for food and an excellent model organism for scientific research (Yan et al., 2014). The quality of genomic data of chicken fulfills the requirements for systematically studying genetic variation. Genetic variation can be studied using the chicken reference genome, including small structural variants, micro, midsize, and large segmental deletions, and insertions in both genic and intergenic regions within the species and closely related species. Studies on the various domestic chicken breeds have identified different sizes of indels associated with phenotypes across the genome (Cui et al., 2006; Dong et al., 2018; Driscoll et al., 2009; Fu et al., 2020; Jin et al., 2016; Kerje et al., 2004; Li et al., 2021; Lin et al., 2021; Liu et al., 2019; Tuanhui Ren et al., 2020; T. Ren et al., 2020; Tang et al., 2011; Wang et al., 2021; X. Wang et al., 2020; Xiangnan Wang et al., 2020; Wei et al., 2020; Yang et al., 2019; Zhou et al., 2010; Zhu et al., 2022). The study of small/midsize and large segmental deletions and insertions in the chicken genome can help to understand their crucial role in the genome size evolution of birds and their association with phenotypic traits. It has been reported that the domestic chicken (*Gallus gallus domesticus*) was domesticated from *Gallus gallus spadiceus*, a subspecies of *Gallus gallus* (red junglefowl) about 8000-9500 years ago (Tixier-Boichard et al., 2011; M. S. Wang et al., 2020). In the *Gallus* genus, red junglefowl has three wild relatives, i.e., *Gallus varius* (green junglefowl), the most ancient lineage, while *Gallus lafayettii* (Ceylon Junglefowl) and *Gallus sonneratii* (grey junglefowl) have a sister relationship with each other and form a sister clade with red junglefowl (Lawal et al., 2020). All four species of the Gallus genus are sexually dimorphic; females are smaller in size and weight (McGowan, P. J. K., G. M. Kirwan, 2020a, 2020b; McGowan, 2020a, 2020b; Morejohn, 1968; Singh et al., 2020). The karyotype and chromosome number (2n=78) of all junglefowl are almost the same, while some variation in genome size has been reported (Okamoto et al., 1988; Piégu et al., 2020) see **Fig. 1**.

**Fig. 1:**
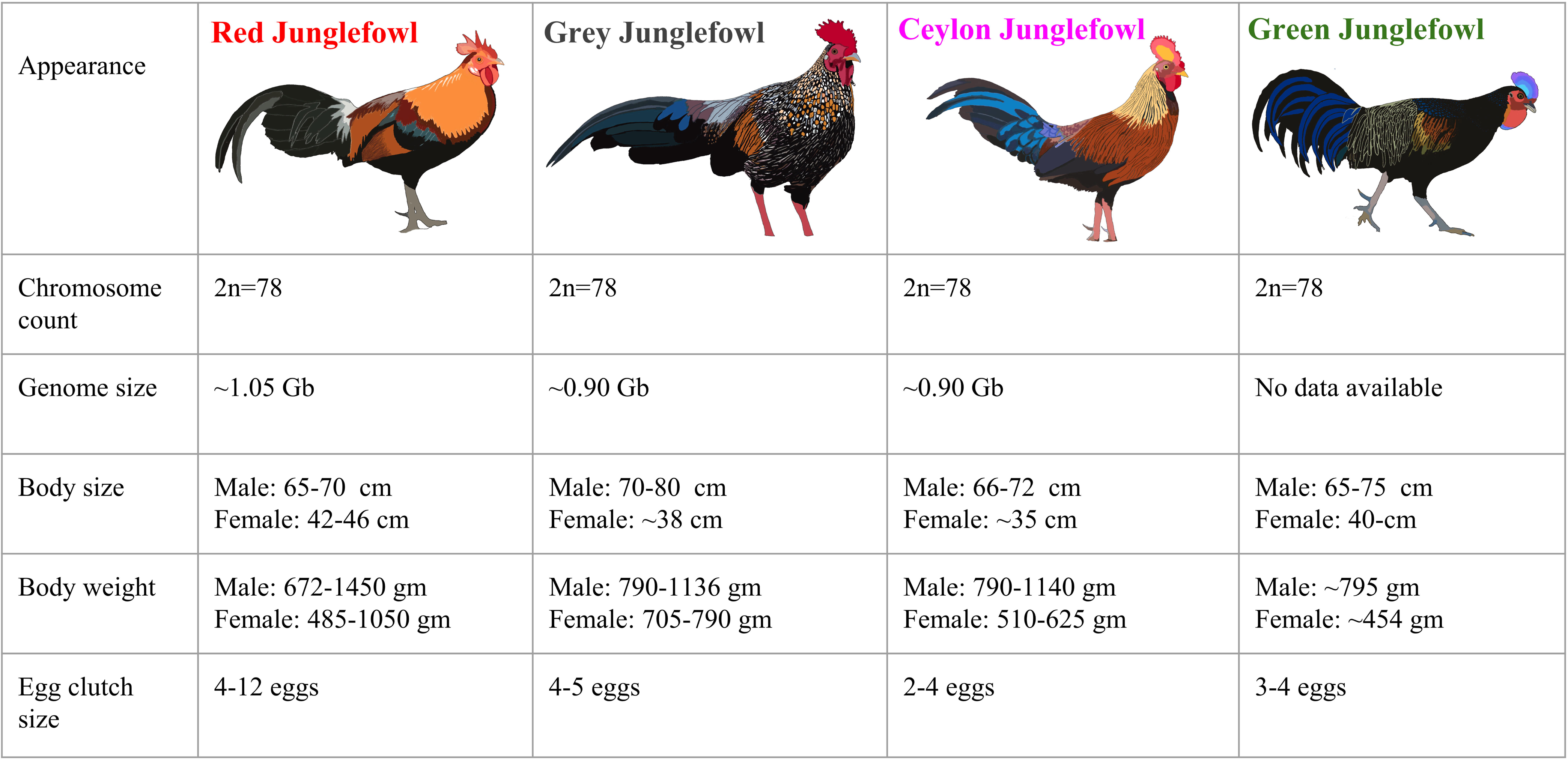
Comparison of the four jungle fowl species of the *Gallus* genus: In the comparison of plumage, *Gallus varius* (green junglefowl) and *Gallus sonneratii* (grey junglefowl) have the most distinct color, while *Gallus gallus* (red junglefowl) and *Gallus lafayettii* (Ceylon junglefowl) resemble each other in form and color (Morejohn, 1968; Singh et al., 2020). The karyotype and chromosome number (2n=78) are the same, while genome size variation has been reported (Okamoto et al., 1988; Piégu et al., 2020). Compared to the cocks, hens are smaller in size and weight in all four species, along with variation in egg clutch size (McGowan, P. J. K., G. M. Kirwan, 2020a, 2020b; McGowan, 2020a, 2020b; Morejohn, 1968; Singh et al., 2020).

Domestication of animals is associated with artificial selection and population bottlenecks, and these factors play an essential role in shaping the genetic variation in domesticated populations (Innan and Kim, 2004). The phenotypic variation occurring in wild species is subject to much weaker selection and less likely to persist. In contrast, in a domestic species, human-mediated artificial selection for a specific trait makes it fixed in a particular population. The evolution of domesticated populations increases genetic differences between domesticated species and their wild ancestors (Andersson and Purugganan, 2022). Previous studies have identified a considerable difference in the number of indels in wild species compared to the domestic populations of sheep (Li et al., 2020; Lv et al., 2022), yak (Wang et al., 2014), dog (Amiri Ghanatsaman et al., 2020), silkworm (Zhang et al., 2015), and rice (Stein et al., 2018). Micro/midsize deletions and insertions have been studied between chicken and phylogenetically more distant species such as Mallard duck, turkey, rock pigeon, and other passerine birds (Brandström and Ellegren, 2007; Kapusta et al., 2017). Even in different breeds of red junglefowl, the genome size variation has been studied to understand the potential role of segmental duplications, deletion, and repeats (Piégu et al., 2020). Compared to the previous studies of micro, midsize deletions and insertions in different breeds of chicken, in this study, we used four closely related species of the *Gallus* genus, which has a divergence time range of 1-4 Mya (Lawal et al., 2020), to identify the midsize (> 1 Kb) deletion events. Studying midsize deletions is challenging because multiple events of microdeletions at the exact location in the genome can give birth to a midsize/large segmental deletion. The role of midsize deletions in shaping genomic architecture via genome size reduction in wild junglefowl species compared to a domestic breed may provide useful information about evolution in the *Gallus* genus. In this study, we aimed to examine the following questions. **1)** Identify midsize deletions (>1 Kb) in wild junglefowl species compared to red junglefowl. **2)** Role of midsize deletion in the genome evolution of 4 closely related species of the *Gallus* genus. **3)** Variation in deletion abundance along the phylogeny of the *Gallus* genus. **4)** Assessment of the evolutionary conservation score of the deleted regions. **5)** Study of candidate genes harboring the deletions and their association with phenotypic traits such as body development and growth, behavioral and plumage color reported in previous studies.

## Materials and Methods

### Data collection, read mapping, and variant calling

To evaluate the genome-wide pattern of midsize deletions (>1 Kb) in the three wild junglefowl species (green junglefowl, Ceylon junglefowl, and grey junglefowl) in comparison to the red junglefowl, we downloaded the publicly available Illumina paired-end resequencing data from Europen Nucleotide Archive (ENA) with >20x coverage. We collected five individuals’ data for each green and Ceylon junglefowl and three for the grey junglefowl. Similarly, we collected data from seven red junglefowl individuals data from two populations to act as controls (for more details, see **Supplementary Table 1**).

We mapped the paired-end raw read Illumina data of 20 individuals to the (Gallus_gallus.GRCg6a) red junglefowl genome assembly with default flags using BWA (Burrows-Wheeler Aligner) v0.7.17-r1188 read mapper (Li and Durbin, 2009). We used the Picard tools (https://github.com/broadinstitute/picard) to add the read group information and remove duplicates from the bam files. We performed the variant calling for all chromosomes except unplaced scaffolds, MT, Z, and W, using two variant callers (bcftools v1.9 (Li and Barrett, 2011) and FreeBayes v1.0.0 (Garrison and Marth, 2012)). To ensure the identification of reliable single nucleotide polymorphisms (SNPs), common SNPs identified by both variant callers were considered using VCFtools v0.1.15 (Danecek et al., 2011), while avoiding false positives. We constructed the maximum likelihood-based phylogenetic tree using SNP data with 1000 bootstraps using SNPhylo (Lee et al., 2014) and visualized it using FigTree v1.4.2 (Rambaut, 2014).

### Identification of genome-wide deletions based on read coverage

We employed a systematic and conservative approach for identifying genome-wide deletions based on read coverage. Firstly, we generated non-overlapping 1Kb windows across the genome using bedtools (Quinlan and Hall, 2010). Subsequently, we calculated the read coverage for each window using the “bedtools coverage” flag for all 20 individuals. Regions with less than five reads in a 1 Kb window (minimum read count criteria) were considered potential midsize deleted regions. After defining the minimum read count criteria as potentially deleted regions, we required all individuals of a particular species to share this putative deletion to be considered further. If the deleted region is continuous and more prolonged than 1 Kb, we used the “bedtools merge’ flag to define the boundaries of deletion regions. To account for the potential false-positive deletion regions caused by overlapping repeat content, we acquired the repeat distribution information for the Gallus_gallus.GRCg6a genome assembly in bed format from the UCSC genome browser, We calculated the coverage of repeat distribution for each 1Kb window. We only considered validated deletion regions with 0% repeat content for further analysis. We excluded the two large regions containing Ns indicating a gap or assembly break in the red junglefowl genome assembly.

### Validation of midsize deletions based on flanking reads

We used short-read Illumina data to confirm the authenticity of midsize deletions from the potential candidate regions. In closely related species, it is possible to rely upon mapped reads to identify segmental deletions. We defined the boundaries of the deleted region and considered a 2 Kb flank region on both sides. In the case of actual deletion, a single read should map on both flanks of the deleted region. We carefully examined such reads that occurred in both flank regions and were positioned at the edge of the deleted region. At the same time, we required that no read should be mapped in the deleted region. To visualize the distributions of deletions across the genome, we generate a circos plot for all three junglefowl species with red junglefowl as a reference using the circlize (v 0.4.15) (Gu et al., 2014).

### phastCons score for deleted regions

We used phastCons to measure the evolutionary conservation score for deleted regions in the chicken genome (Siepel et al., 2005). We downloaded the phastCons score from https://hgdownload.soe.ucsc.edu/goldenPath/galGal6/phastCons77way/ and converted it into the bed format. After getting the phastCons conservation score for each deleted region, we plotted it along the length of the deleted regions using R (Ihaka and Gentleman, 1996) to evaluate the conservation status of the deleted regions in red junglefowl.

### Gene enrichment analysis

We discovered multiple >1 Kb deletions in intronic, intergenic, and exonic (long non-coding RNAs) regions, which were verified by manually inspecting the mapped reads. We considered the genes with intronic deletions and those located within ∼50 Kb downstream from the boundary of deletions as those potentially affected. We prepared the list of ENSEMBL gene ids for these genes for all three junglefowl species and used the ShinyGO 0.76.2 server (Ge et al., 2020) for the gene enrichment analysis.

## Results

### Distribution of midsize deletions in the *Gallus* genus

To identify the >1 Kb deletions, we examined four junglefowl species: red junglefowl, grey junglefowl, Ceylon junglefowl, and green junglefowl. We utilized the whole genome resequencing data of seven individuals of red junglefowl, three grey junglefowl, five green junglefowl, and Ceylon junglefowl. We constructed a phylogenetic tree based on autosomal single nucleotide polymorphisms (SNPs) data, which suggest that the green junglefowl diverge first, followed by Ceylon, grey, and red junglefowl, respectively (**Fig. 2A**). The phylogenetic tree is supported by the previously published genome-wide phylogeny of the *Gallus* genus (Lawal et al., 2020). We observed a trend of an increasing number of midsize deletions with increasing phylogenetic distance from red junglefowl, which suggests a linear accumulation of deletions. Notably, these deletions do not result from sequence divergence, as the same mapped reads span both sides of the deletion. Among the junglefowl species, green junglefowl has the highest number (77) of deletions, followed by the Ceylon junglefowl (41). In contrast, grey junglefowl has the lowest number (12) (**Fig. 2B, Fig. S1,** and **Supplementary Table 2-4**). Three deleted regions were common between Ceylon and grey junglefowl, while two were common in Ceylon and green junglefowl. No commonly deleted regions could be identified between green and grey junglefowl and among all three species. Additionally, we identified a region commonly lacking coverage in all junglefowl species resulting from an assembly gap in the *Gallus_gallus*.GRCg6a genome assembly (**Supplementary Table 5**). Midsize deletions were distributed throughout the genome in all three junglefowl species occurring in exons of some lncRNA’s (long non-coding RNA), intergenic and intronic regions. The length of deletions identified genome-wide varied, ranging from 1 Kb to 2.8 Kb in grey junglefowl, 1 Kb to 5 Kb in Ceylon junglefowl, and 1 Kb to 10 Kb in green junglefowl (**Fig. S2, Supplementary Table 2**). A total length of 23 Kb in grey, 92.1 Kb in Ceylon, and 178.9 Kb in green junglefowl have been deleted compared to red junglefowl.

**Fig. 2:**
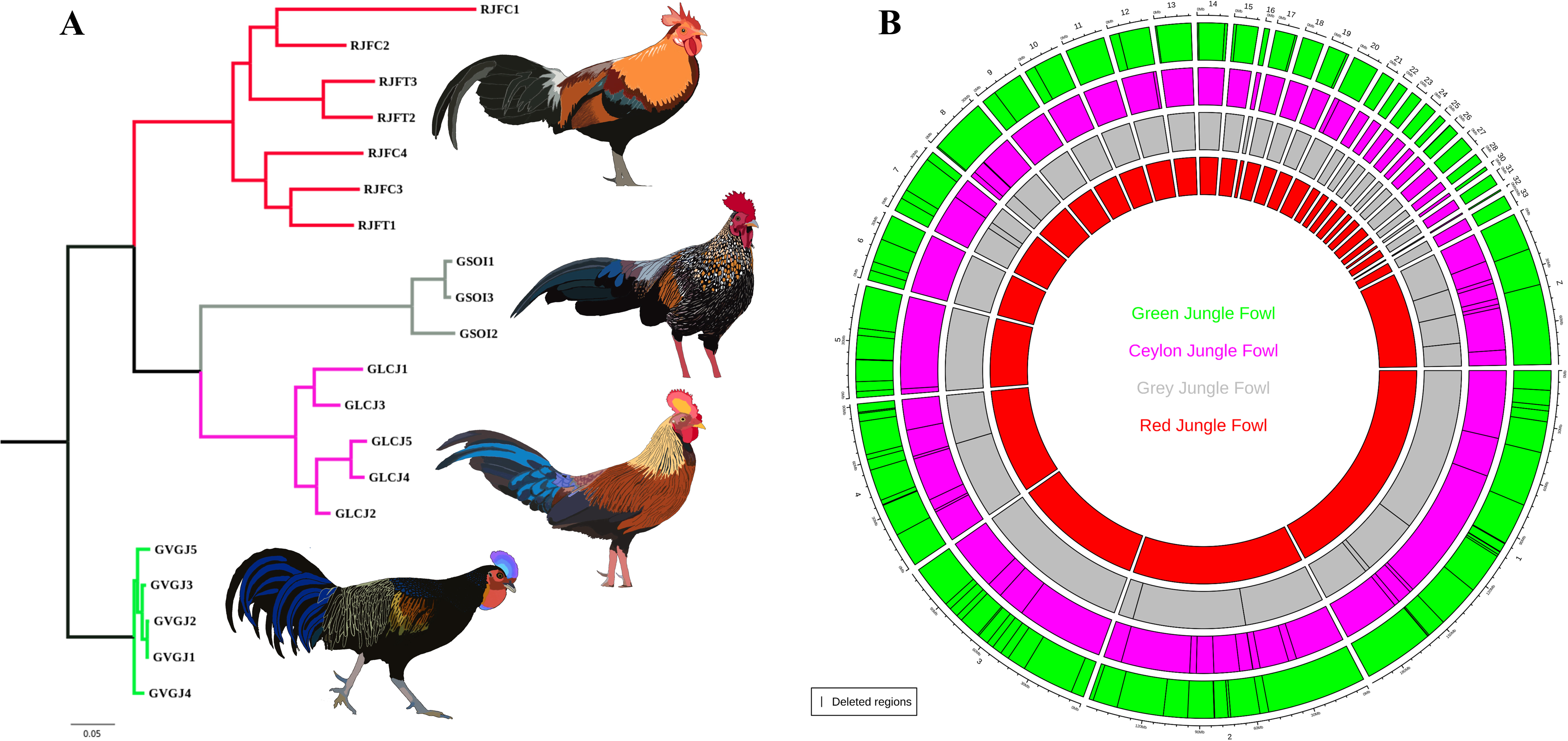
Phylogenetic relationships and distribution of deletions in *Gallus* genus: **A**. Genome- wide phylogenetic tree of four junglefowl species of *Gallus* genus. Species-specific colors are assigned in the phylogeny (such as red for junglefowl, grey for grey junglefowl, magenta for Ceylon junglefowl, and green for green junglefowl). **B.** Genome-wide deletion pattern across three junglefowl species compared to the red junglefowl species. The circlize plot show circles for the particular species and the location of deleted regions shown by the inverted black lines.

### Midsize deletions in grey junglefowl

Four of the twelve midsize deletions detected in the grey junglefowl species are located in functionally important genes, such as *DIAPH3, GLIS3, ADAMTS19,* and *SCN1A,* which harbor deletions in the intronic and upstream regions. Diaphanous homolog 3 (*DIAPH3*) is located on chromosome 1 in the red junglefowl genome assembly and contains 23 exons encoding a protein of 1025 amino acids. *DIAPH3* harbors >2.5 Kb deletion in the 17^th^ intron verified using mapped reads (**Fig. 3A**). We assessed the mean coverage of the same deleted region of all four junglefowl species for further investigation (**Fig. S3**). Clear evidence of deletion, specific to the grey junglefowl, was observed (**Fig. 3B**). Notably, the deleted region within *DIAPH3* exhibits a high phastCons conservation score in the red junglefowl genome assembly. (**Fig. 3C**). *GLIS3* and *ADAMTS19* genes also have a deletion in their second and the twentieth intron, respectively (**Fig. S4, 5**). The size of deletions in these two genes is 2.4 Kb and 1.1 Kb, respectively (**Supplementary Table 2**). Furthermore, an intergenic deletion, with a size of 1.3 Kb, is located at ∼22Kb upstream of the *SCN1A* gene (**Fig. S6**). Detailed information on remaining deletions identified in grey junglefowl is provided in **Supplementary Table 2**.

**Fig. 3:**
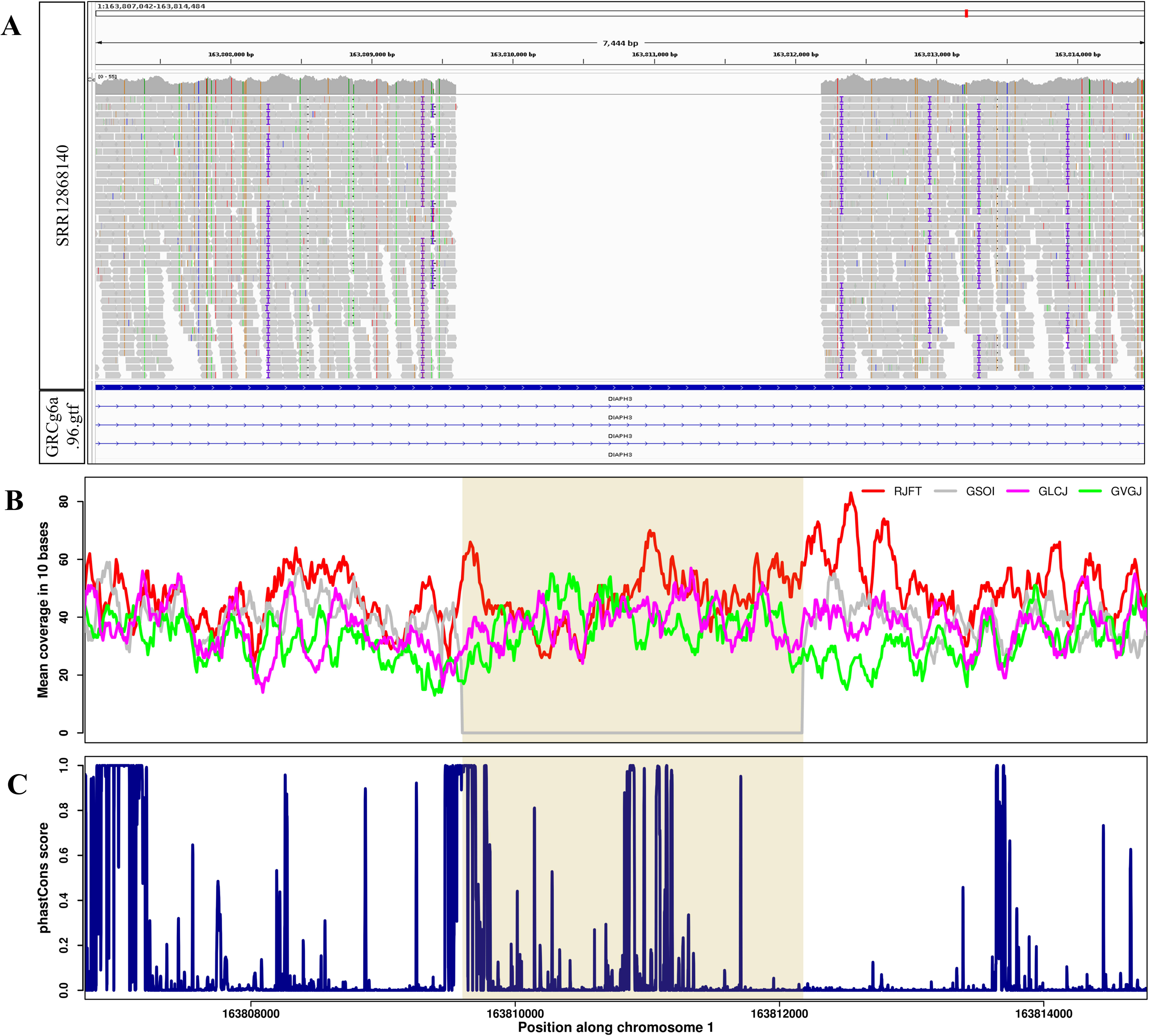
Midsize deletion in grey junglefowl: **A.** IGV screenshot of *DIAPH3* intronic region deletion located at Chr 1 in grey junglefowl (SRR12868140) species. Grey horizontal rectangular boxes represent the reads mapped with the Gallus_gallus.GRCg6a genome assembly and the white space represent the deleted region. The General Feature Format (GFF/GTF) is loaded at the bottom. **B.** The mean coverage per ten base windows of four species with the deleted region is highlighted using transparent tan color. **C.** The phastCons score of the Gallus_gallus.GRCg6a genome assembly of deleted and adjacent regions is represented in dark blue.

### Midsize deletions in Ceylon junglefowl

In our study of Ceylon junglefowl, we identified 13 midsize deletions within the intronic region of different genes. Additionally, we identified 26 deletions in intergenic regions. Interestingly two deletions were found to overlap exons, resulting in the removal of exon2 from the ENSGALG00000052188 (lncRNA) gene and exon5 from the ENSGALG00000049888 (lncRNA) gene (**Supplementary Table 3**). Out of 13 intronic deletions, we focused on three regions within *KCNH8*, *GRIK2*, and *NDST4* genes based on their conservation score and known functions. The *KCNH8* gene is 16 exonic and located on chromosome 2 in the chicken genome. We observed the deletion of approximately 3.8 Kb within the second intron of *KCNH8,* which has been verified by mapped reads in Ceylon junglefowl (**Fig. 4A**). The evaluation of mean coverage in 10bp windows of this deleted region for all four junglefowl species further supports the presence of this midsize deletion in Ceylon junglefowl (**Fig. 4B** and **Fig. S7**). Notably, this deleted region has a high phastCons conservation score in red junglefowl genome. (**Fig. 4C**). We detected midsize deletion in the remaining two genes, GRIK2 and NDST4, in their sixth and second intron with a size of 1.3 Kb and 1.4 Kb, respectively **(Fig. S8, 9** and **Supplementary Table 3**). Three intergenic deletions were located approximately 35 Kb, 49 Kb, and 16 Kb upstream from *SQLE*, and a paralogous copy of *COL11A1* and *LMCD1* genes (**Fig. S10-12**). Detailed information on remaining deletions is provided in **Supplementary Table 3**.

**Fig. 4:**
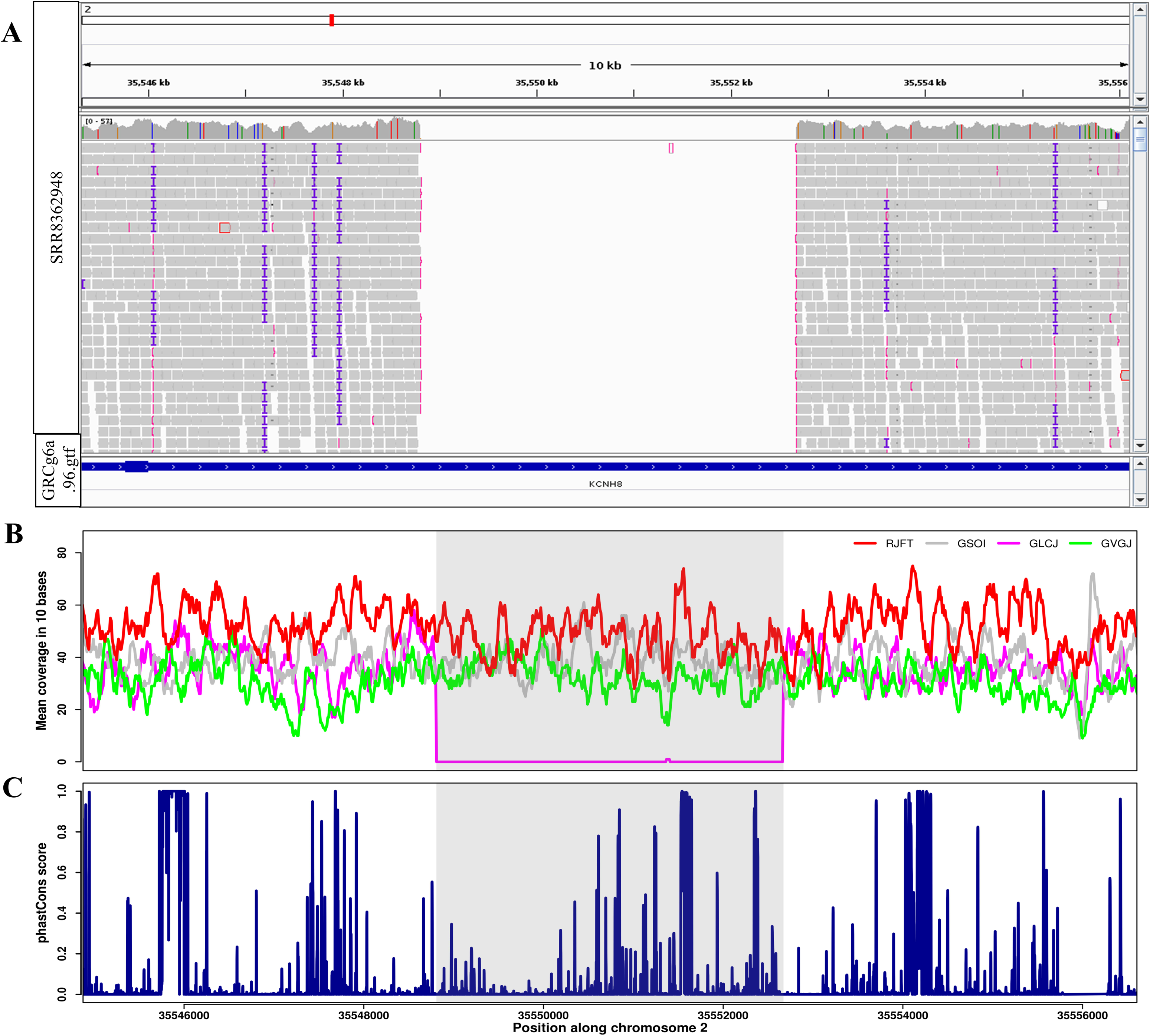
Midsize deletion in Ceylon junglefowl: **A.** IGV screenshot of *KCNH8* intronic region deletion located at Chr 2 in Ceylon junglefowl (SRR8362948) species. Grey horizontal rectangular boxes represent the reads mapped with the Gallus_gallus.GRCg6a genome assembly and the white space represent the deleted region. **B.** The mean coverage per ten base windows of four species with the deleted region is highlighted using transparent tan color. **C.** The phastCons score of the Gallus_gallus.GRCg6a genome assembly of deleted and adjacent regions is represented in dark blue.

### Midsize deletions in green junglefowl

In our study of green junglefowl, we found 32 deletions in the intronic region of different genes, 41 deletions in the intergenic region, and 4 deletions in the exon of different lncRNAs **(Supplementary Table 4**). Out of 77 deletions, we focused on 9 (8 intronic and one intergenic), which are functionally important and have high conservation scores. Notably, a deletion of 1.4 Kb length was observed in the third intron of the *CACNA2D1* gene, located on chromosome 1 in the red junglefowl genome (**Fig. 5A**). When comparing the read coverage in this region across the remaining three junglefowl species we found coverage of at least five reads compared to zero reads in the green junglefowl (**Fig. 5B and Fig. S13**). The red junglefowl genome has high conservation for some parts of this deleted region (**Fig. 4C**). The *KIF26B*, *VIP*, *CCSER1*, *SGCZ*, *LRP1B*, *SHISA9*, and *TBC1D22B* genes also have deletions in their introns (**Fig. S14-20**). For the estimated size of the deletions in these genes, see **Supplementary Table 4**. One intergenic deletion of size 1.4 Kb is located at 600 bp upstream of the *LRPAP1* gene (**Fig. S21**). The information on the remaining deletions reported in green junglefowl is given in **Supplementary Table 4**.

**Fig. 5:**
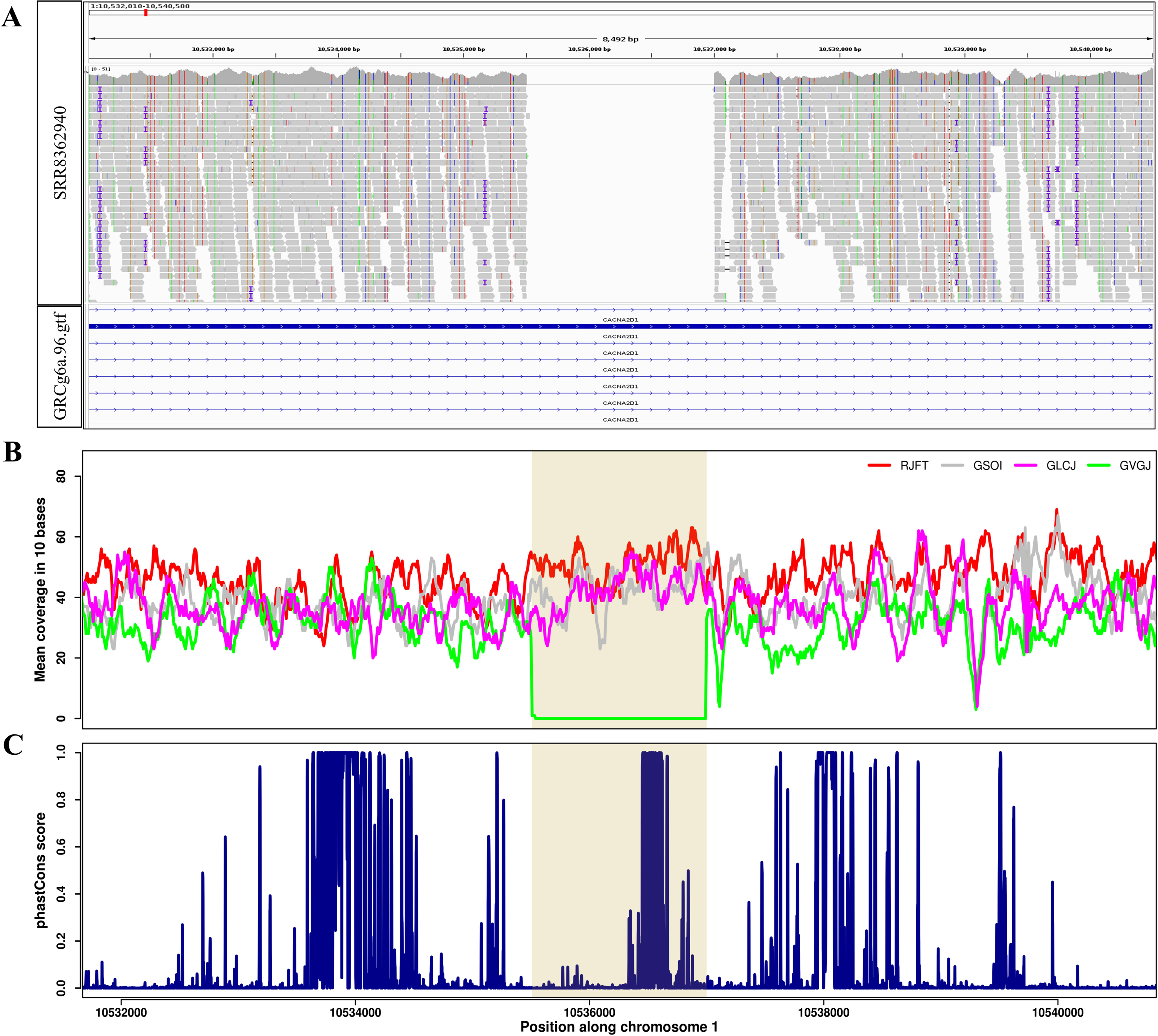
Midsize deletion in green junglefowl: **A.** IGV screenshot of *CACNA2D1* intronic region deletion located at Chr 1 in green junglefowl (SRR8362940) species. Grey horizontal rectangular boxes represent the reads mapped with the Gallus_gallus.GRCg6a genome assembly and the white space represent the deleted region. The General Feature Format (GFF/GTF) is loaded at the bottom. **B.** The mean coverage per ten base windows of four species with the deleted region is highlighted using transparent tan color. **C.** The phastCons score of the Gallus_gallus.GRCg6a genome assembly of deleted and adjacent regions is represented in dark blue.

### Gene enrichment analysis

The gene enrichment analysis of candidate genes identified in grey junglefowl species shows enrichment for the SOSS complex, Voltage-gated sodium channel complex, sodium channel complex, and multiple micro RNAs targeted gene pathways (**Fig. S22** and **Supplementary Table 6**). In Ceylon junglefowl, various gga-mir target gene pathways and biosynthesis pathways show the enrichment in GO molecular function, curated PANTHER, and KEGG pathway analysis (**Fig. S23** and **Supplementary Table 6**). In the case of green junglefowl, various gga-mir target gene pathways, PDZ domain binding, negative regulation of actin filament bundle assembly, and system process pathways are enriched (**Fig. S24 and Supplementary Table 6**).

## Discussion

The main objective of this study is to gain a comprehensive understanding of the distribution and prevalence of midsize deletions (≥1 Kb) throughout the genomes of three distinct wild junglefowl species (grey, Ceylon, and green). The red junglefowl species is used as a reference point to identify deleted regions. Species within the *Gallus* genus are phylogenetically closely related, with a divergence time of 1-4 million years ago (Lawal et al., 2020), making them an ideal system for studying segmental deletions. While previous research has investigated insertions and deletions in more distantly related species like duck, turkey, rock pigeon, and other passerine birds in comparison to chicken (Kapusta et al., 2017), an investigation specifically targeting midsize deletions within wild junglefowl species has yet to be conducted. Consequently, our study aims to bridge this gap in knowledge. The rationale behind prioritizing midsize deletions larger than 1 Kb is rooted in comprehending their impact on DNA loss, genome size evolution, and their distribution patterns within the genome. Given their size and genomic positioning, midsize deletions can significantly influence the fitness of a species, as they possess the potential to be deleterious. In nature, any specific trait resulting from an insertion or deletion undergoes selection only if it enhances the fitness of the species. Therefore, the midsize deletions currently fixed or present in a high frequency in wild junglefowl species may offer valuable insights into their functional consequences.

The distribution and cumulative length of midsize deletions observed in three junglefowl species provide insights into the DNA loss process during the evolution of the *Gallus* genus. Among the species studied, the green junglefowl, being the farthest in terms of phylogeny based on SNP data, exhibited the highest number of deletions, followed by Ceylon and grey junglefowl in decreasing order. This increasing trend in the number of midsize deletions with phylogenetic distance supports the concept of linear accumulation of such deletions, as discussed in the mutational equilibrium model (Petrov, 2002b). In our study, we found approximately 23 Kb of DNA loss in grey junglefowl and approximately 92 Kb in Ceylon junglefowl, which accounts for 0.0022% (23000 * 100 / 1050000000(the genome size of red junglefowl)) and 0.0087% (92000 * 100 / 1050000000) of genome loss, respectively. Considering a phylogenetic distance of 2.6-2.9 My from red junglefowl, this suggests a rate of DNA loss during *Gallus* genus evolution of 7.93-8.84 Kb/My in grey junglefowl and 31.72-35.38 Kb/My in Ceylon junglefowl. In the case of green junglefowl, approximately 178 Kb of DNA loss accounts for 0.017% (178 * 100 / 1050000000) of the genome loss, with a rate of approximately 44 Kb/My of DNA loss. Previous research has reported that small/microdeletions contribute to 1% of DNA loss in chickens compared to the Galloanserae and Neoaves lineages, while no data is available for midsize deletions (Kapusta et al., 2017). The relatively lower percentage of DNA loss attributed to midsize deletions in our study of the *Gallus* genus could be due to our stringent criteria and the shallow phylogenetic distances between the junglefowl species.

The *Gallus* genus has four species, each displaying sexual dimorphism and distinct plumage colors. Plumage color, a polygenic trait, can arise from gene-gene epistatic interactions, regulatory genes, or the influence of multiple coding genes (Davoodi et al., 2022). Studies examining different chicken breeds have identified associations between upstream and intergenic deletions and variations in plumage color (Bortoluzzi et al., 2020; Gunnarsson et al., 2011; Zhu et al., 2022), implying a potential role of deletions in determining plumage color. The egg clutch size, body size, and weight are comparable between the species of the *Gallus* genus. The genes associated with body development and growth traits, adaption against environmental conditions, and plumage color have essential roles in survival and evolution in the wild. Therefore, we focused on deleted regions in the intronic and upstream regions of such genes. Since functional studies specific to grey, Ceylon, and green junglefowl are not available, we rely on previous studies primarily conducted on domestic chickens (refer to **Table 1**). The associations between candidate genes and various phenotypic traits reported in these previous studies can provide insights into the impact of midsize deletions in wild junglefowl. The domestication process has led to various changes in traits among domesticated species compared to their wild counterparts (Diamond, 2002). Specific traits such as variations in plumage color, reduction in bone mass and cranial volume, as well as behavioral traits like reduced anxiety, fear, and aggression, have been observed in domesticated animals (Price, 1999; Trut, Oskina and Kharlamova, 2009).

**Table 1:**
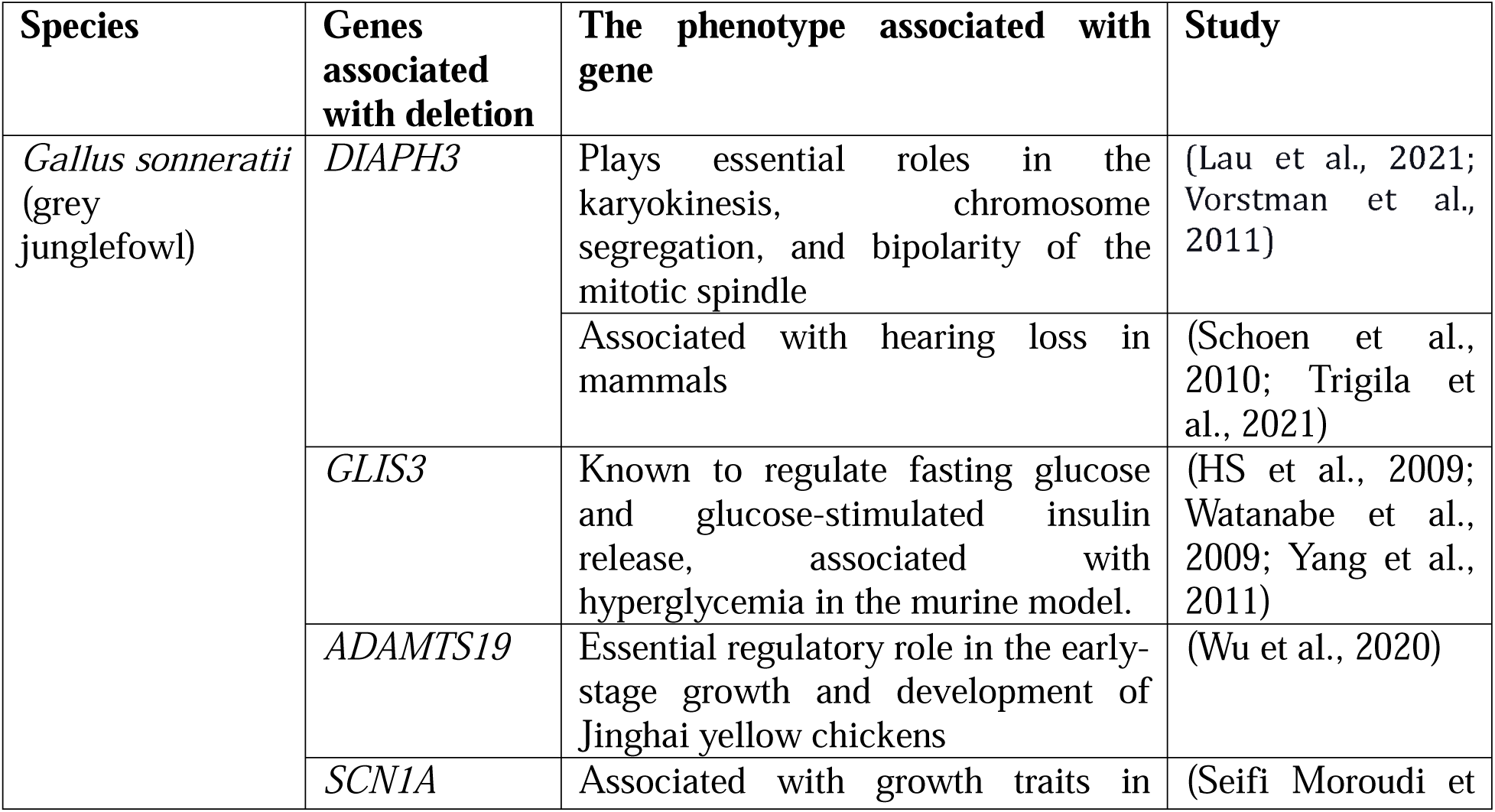
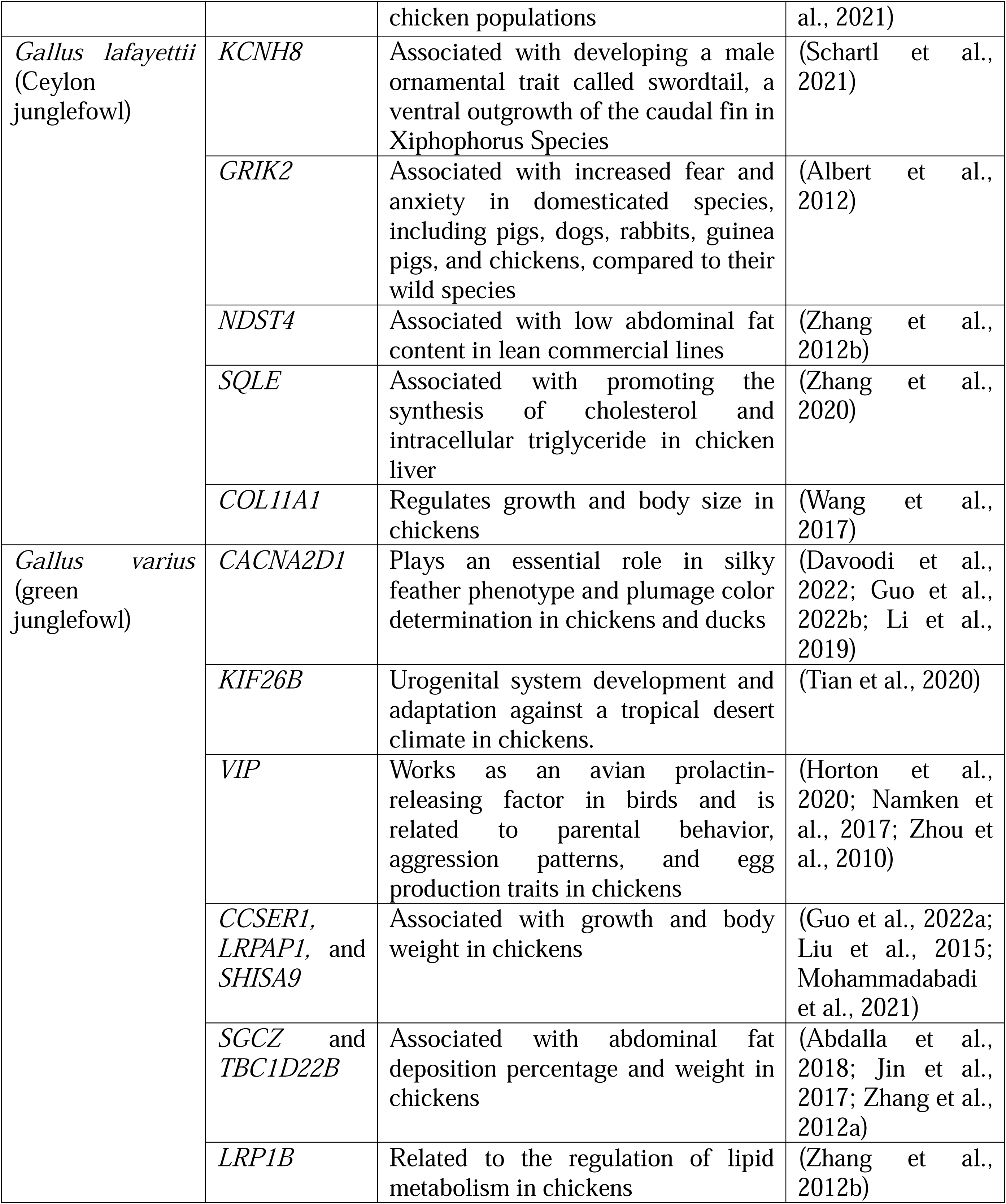
List of genes harboring deletions in their intronic region and upstream region identified in wild relatives of the red junglefowl.

In grey junglefowl, midsize deletions found in the intronic regions of *DIAPH3*, *GLIS3*, and *ADAMATS19*, as well as an upstream deletion in *SCN1A*, may have associations with cytoskeleton maintenance, neural development, enhancement of hearing capacity, and body development and growth traits (refer to **Table 1)**. Ceylon junglefowl exhibits a more vibrant orange-red body plumage color, highly exaggerated wattles, and a red-colored comb with a yellow patch (Morejohn, 1968; Silva and Rajapaksha, 2005). The deletion in the second intron of the *KCNH8* gene could potentially impact body and comb traits, as it is associated with the development of a male ornamental trait known as the swordtail in swordtail fish species *Xiphophorus helleri* (Schartl et al., 2021). Midsize deletions in the intronic regions of *GRIK2* may be linked to aggressive behavior, fear, and anxiety, as increased expression of the *GRIK2* gene has been associated with heightened fear and anxiety in domesticated species such as pigs, dogs, rabbits, guinea pigs, and chickens compared to their wild counterparts (Albert et al., 2012). The observation of aggressive behavior in junglefowl species is often attributed to the maintenance of social hierarchy and competition for limited resources within small groups of birds (Queiroz and Cromberg, 2006). Midsize deletion in the second intron of *NDST4* and upstream direction of *SQLE* and a paralogous copy *COL11A1* may be associated with lean body type, low abdominal fat and cholesterol content, and growth and body size **(Table1)**. The wild junglefowl’s abdominal fat and cholesterol content is reportedly low compared to domestic chickens (Rahayu et al., 2008). Green junglefowl, which diverged approximately 4 million years ago from the common ancestor of wild junglefowl species, exhibits distinct phenotypic characteristics, including dark center orange and yellow feathers and a vibrant blue and pink comb (Lawal et al., 2020). The striking plumage coloration and comb appearance in green junglefowl might be influenced by a 1.4 Kb deletion within the third intron of *CACNA2D1*, a gene known to play a significant role in silky feather phenotype and plumage color determination in chickens and ducks (Davoodi et al., 2022; Guo et al., 2022b; Li et al., 2019). Deletions in the intronic regions of *KIF26B*, *VIP*, *CCSER1*, *LRPAP1*, *SGCZ*, *TBC1D22B*, and *LRP1B* might be associated with adaptation, aggression patterns, egg production, growth, and body weight, abdominal fat deposition percentage, and weight traits **(Table1).**

Due to the relatively low divergence time of 1-4, Mya, the short-read data of the other three species can be mapped to the red junglefowl genome assembly with a high mapping quality and provides a well-suited system to study midsize deletion in the *Gallus* genus. Our methodology followed stringent criteria to address challenges posed by repeat regions, polymorphic deletions (where deletion is supported by three out of five individuals, for example), assembly gaps in the red junglefowl genome, and spurious or low-quality mapped reads within deleted regions. Such excluded regions from our study could be further explored with the availability of high-quality genome assemblies and long-read data for the other three junglefowl species (refer to **Supplementary Fig. 25-28)**. We found deletions in the exonic regions of long non-coding RNA genes annotated in the chicken genome. Comparison with the duck genome indicated the absence of protein-coding genes in these regions. Hence, further improvements in genome annotation among *Gallus* species are required. Due to the unavailability of high-quality genomes of the other three junglefowl species, we cannot study chromosomal rearrangement events compared to domestic chickens. Such rearrangements between wild and domesticated species have recently been identified using high-quality soybean genomes (Xie et al., 2019). However, conducting comparative and functional studies on grey, Ceylon, and green junglefowl can provide valuable insights into the *Gallus* genus’s evolutionary trajectory and survival strategies. Large segmental deletions pose challenges when using short-read data, as they can be generated by multiple small deletion events, making them challenging to study. Therefore, including long-read data can be advantageous for investigating large segmental deletions. Additionally, the genome-wide alignment approach and advanced genomic alignment tools can be employed to comprehensively study small, midsize, and large segmental deletions.

## Conclusion

Our study has revealed the presence of midsize deletions (>1 Kb) in the genomes of grey, Ceylon, and green junglefowl species compared to the red junglefowl. These deletions are predominantly located in intergenic and intronic regions. Notably, we did identify a few midsize deletions within the exonic regions of long non-coding RNA genes annotated in chicken. Deletions in protein-coding regions may be selectively disadvantageous and thus eliminated through natural selection in the wild junglefowl species. The lengths of midsize deletions were comparable among the three junglefowl species. In contrast, the number of deletions increased with greater phylogenetic distance, with green junglefowl exhibiting the highest number of deletions due to its greater evolutionary divergence. Most midsize deletions occurred independently in each junglefowl species, although a few regions exhibited common deletions between certain species. The genes that have deletions in their intergenic and intronic regions are known to be involved in various phenotypic traits, including plumage color, body development and growth, meat and egg production, adaptation to pathogens, and survival in extreme environmental conditions in domestic chicken. Further investigation of these midsize deletions in grey, Ceylon, and green junglefowl species may provide valuable insights into their functional implications and association with essential genes.

## Acknowledgment

We thank the University Grants Commission for supporting AS with a Ph.D. scholarship and the Council of Scientific & Industrial Research for Ph.D. fellowship to S.S.S. The Department of Biotechnology, Ministry of Science and Technology, India (Grant no. BT/11/IYBA/2018/03) and Science and Engineering Research Board (Grant no. ECR/2017/001430) provided funds used to purchase computational resources (i.e., Har Gobind Khorana Computational Biology cluster) used.

## Appendix

Supplementary 1. Supplementary tables

Supplementary 2. Supplementary figures

## Availability of data

Scripts and data are available at: https://github.com/Ashu2195/Midsize_deletion_in_Gallus_genus and https://doi.org/10.17632/k7t62v5frv.1.

## Author contributions

A.S. and S.S.S. wrote the manuscript under the supervision of N.V. A.S., and S.S.S. analyzed the data with inputs from N.V. All authors reviewed the manuscript.

## Notes

https://doi.org/10.17632/k7t62v5frv.1

### Competing Interest Statement

The authors have declared no competing interest.

