## Supplementary Figures for "Hundreds of independent midsize deletions mediate DNA loss in wild relatives of Red Jungle Fowl"

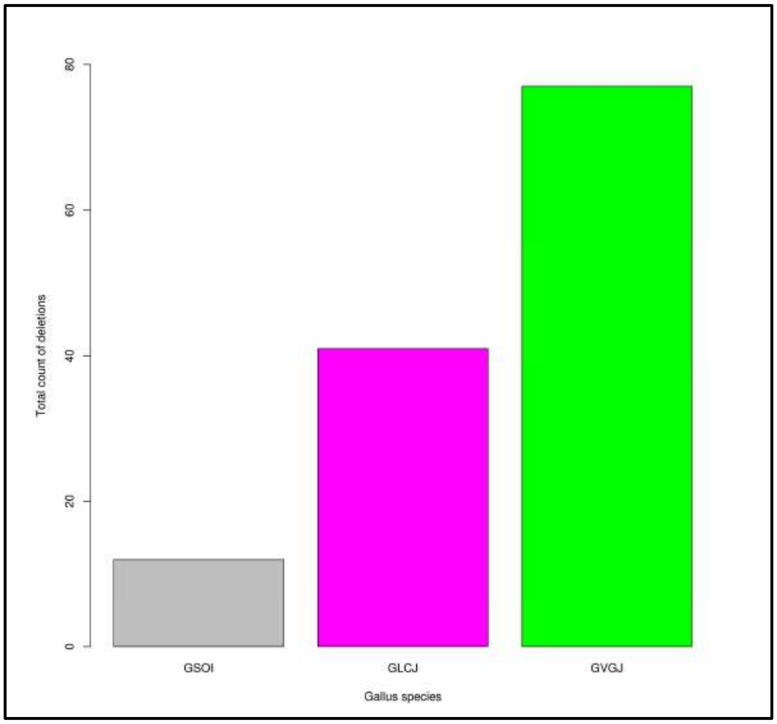


**Fig. S1**: Bar plot of the total number of midsize deletions identified in, grey junglefowl (GSOI), Ceylon junglefowl (GLCJ) and green junglefowl (GVGJ).

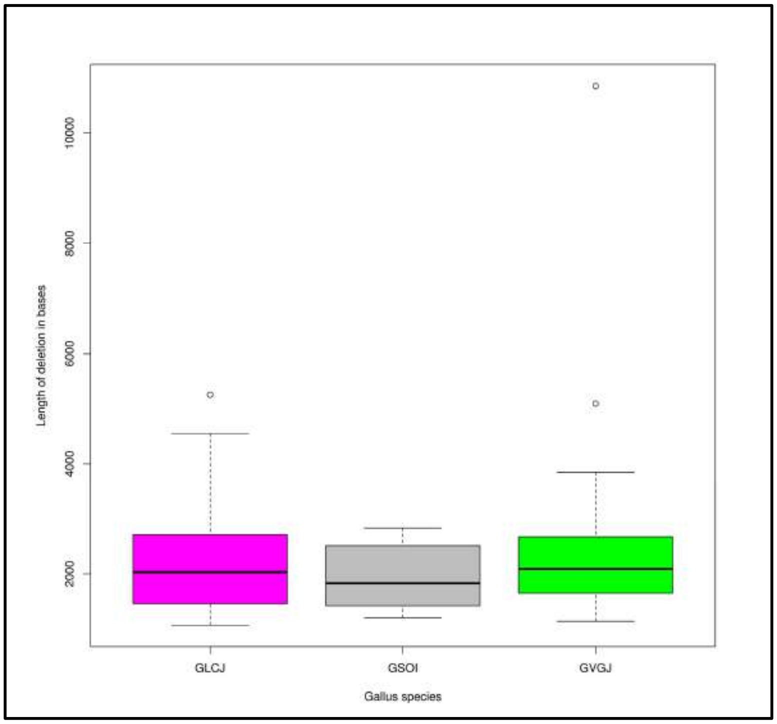


**Fig. S2:** Boxplot of the length distribution of deletions identified in, grey junglefowl (GSOI), Ceylon junglefowl (GLCJ) and green junglefowl (GVGJ).

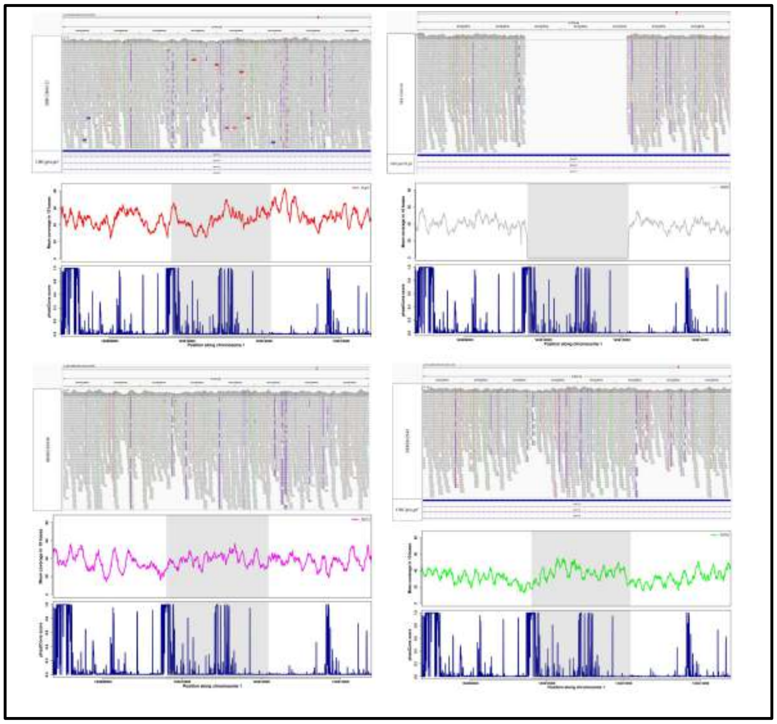

**Fig. S3:** IGV screenshot, mean coverage of 10 bases per window, and phastCons score of the Galgal6 genome assembly of deleted and adjacent regions for four junglefowl species, red junglefowl (SRR12868132), grey junglefowl (SRR12868140), Ceylon junglefowl (SRR8362948), and green junglefowl (SRR8362940) respectively at position 1:163807042-163814484 intronic region of *DIAPH3* gene is shown. Grey horizontal rectangular boxes represent the reads mapped to the Galgal6 genome assembly, and the white space represents the deleted region in grey junglefowl species. The General Feature Format (GFF/GTF) is loaded at the bottom.

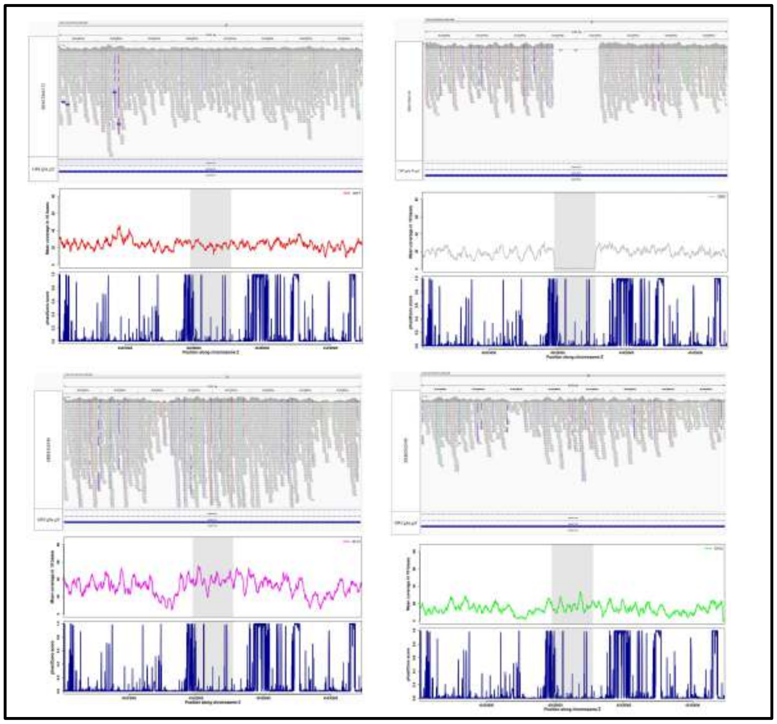

**Fig. S4:** IGV screenshot, mean coverage of 10 bases per window, and phastCons score of the Galgal6 genome assembly of deleted and adjacent regions for four junglefowl species, red junglefowl (SRR12868132), grey junglefowl (SRR12868140), Ceylon junglefowl (SRR8362948), and green junglefowl (SRR8362940) respectively at position Z:45422400-45430600 intronic region of *ADAMTS19* gene is shown. Grey horizontal rectangular boxes represent the reads mapped to the Galgal6 genome assembly, and the white space represents the deleted region in grey junglefowl species. The General Feature Format (GFF/GTF) is loaded at the bottom.

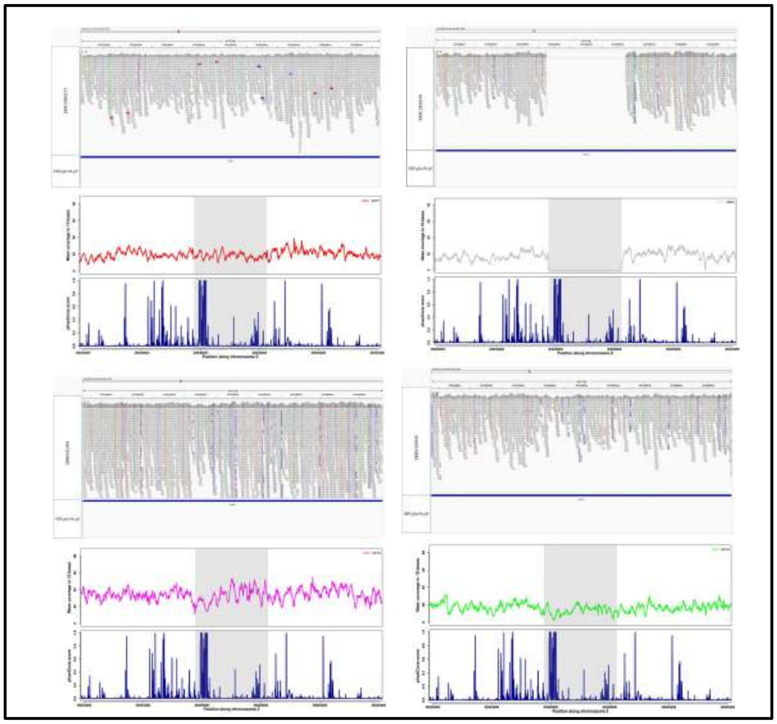

**Fig. S5:** IGV screenshot, mean coverage of 10 bases per window, and phastCons score of the Galgal6 genome assembly of deleted and adjacent regions for four junglefowl species, red junglefowl (SRR12868132), grey junglefowl (SRR12868140), Ceylon junglefowl (SRR8362948), and green junglefowl (SRR8362940) respectively at position Z:26932270-26941740 intronic region of *GLIS3* gene is shown. Grey horizontal rectangular boxes represent the reads mapped to the Galgal6 genome assembly, and the white space represents the deleted region in grey junglefowl species. The General Feature Format (GFF/GTF) is loaded at the bottom.

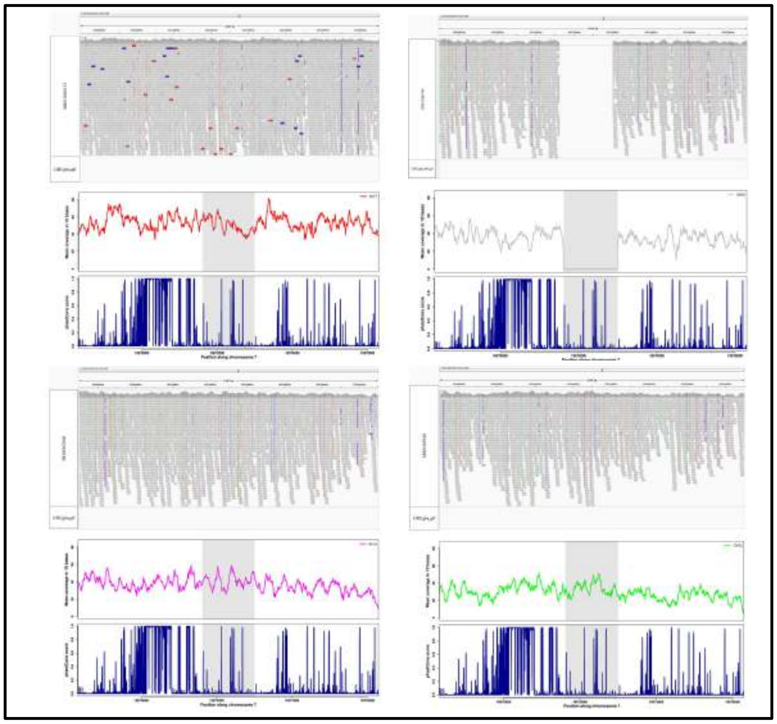

**Fig. S6:** IGV screenshot, mean coverage of 10 bases per window, and phastCons score of the Galgal6 genome assembly of deleted and adjacent regions for four junglefowl species, red junglefowl (SRR12868132), grey junglefowl (SRR12868140), Ceylon junglefowl (SRR8362948), and green junglefowl (SRR8362940) respectively at position 7:19568620-19576000 located in an intergenic region and 22Kb adjacent to *SCN1A* gene are shown. Grey horizontal rectangular boxes represent the reads mapped to the Galgal6 genome assembly, and the white space represents the deleted region in grey junglefowl species. The General Feature Format (GFF/GTF) is loaded at the bottom.

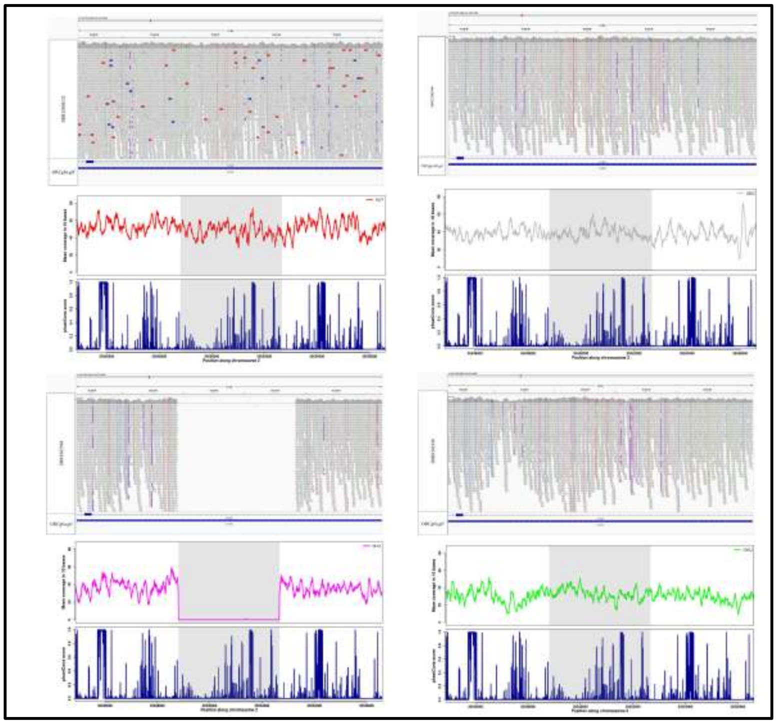

**Fig. S7:** IGV screenshot, mean coverage of 10 bases per window, and phastCons score of the Galgal6 genome assembly of deleted and adjacent regions for four junglefowl species, red junglefowl (SRR12868132), grey junglefowl (SRR12868140), Ceylon junglefowl (SRR8362948), and green junglefowl (SRR8362940) respectively at position 2:35545310-35556170 intronic region of *KCNH8* gene is shown. Grey horizontal rectangular boxes represent the reads mapped to the Galgal6 genome assembly, and the white space represents the deleted region in Ceylon junglefowl species. The General Feature Format (GFF/GTF) is loaded at the bottom.

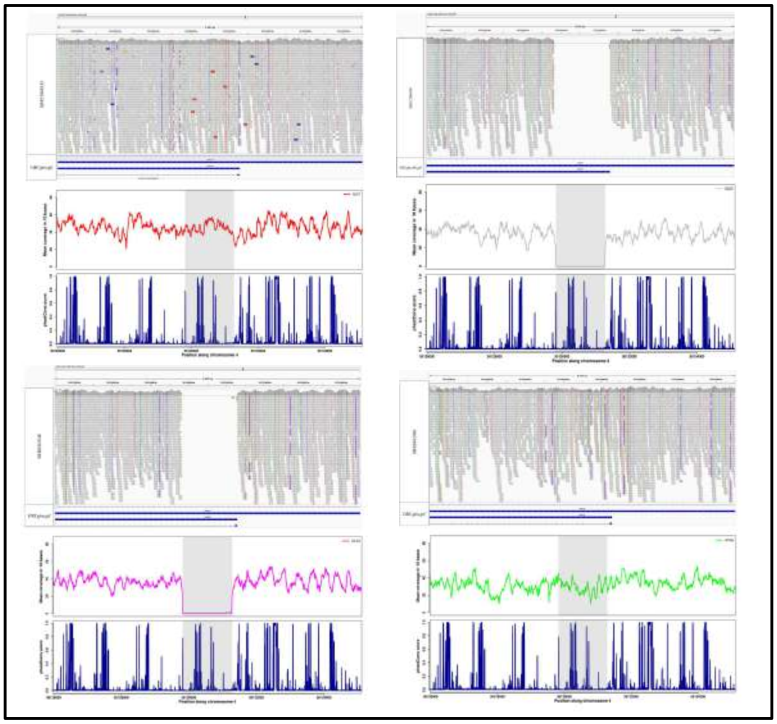

**Fig. S8:** IGV screenshot, mean coverage of 10 bases per window, and phastCons score of the Galgal6 genome assembly of deleted and adjacent regions for four junglefowl species, red junglefowl (SRR12868132), grey junglefowl (SRR12868140), Ceylon junglefowl (SRR8362948), and green junglefowl (SRR8362940) respectively at position 4:56126320-56134780 intronic region of *NDST4* gene is shown. Grey horizontal rectangular boxes represent the reads mapped to the Galgal6 genome assembly, and the white space represents the deleted region in grey junglefowl and Ceylon junglefowl species. The General Feature Format (GFF/GTF) is loaded at the bottom.

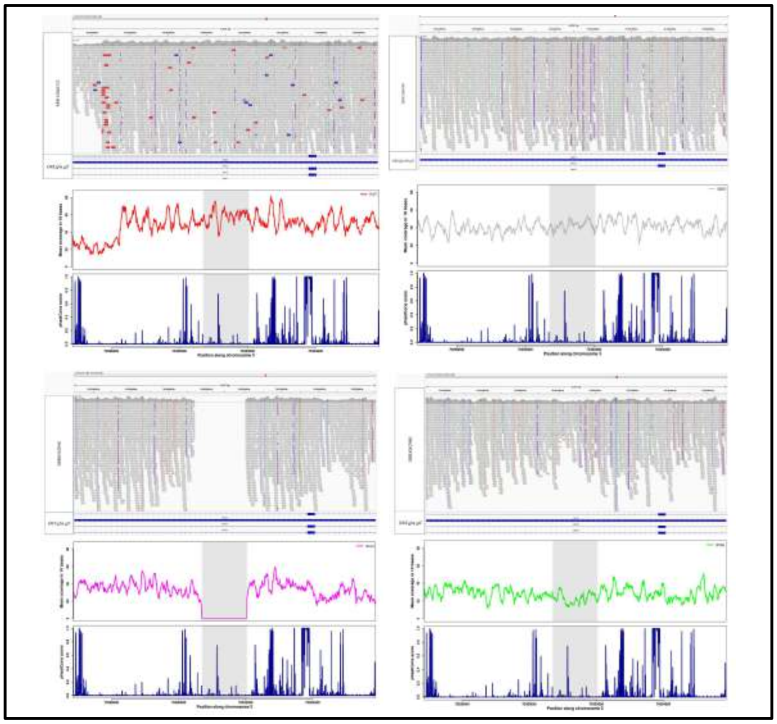

**Fig. S9:** IGV screenshot, mean coverage of 10 bases per window, and phastCons score of the Galgal6 genome assembly of deleted and adjacent regions for four junglefowl species, red junglefowl (SRR12868132), grey junglefowl (SRR12868140), Ceylon junglefowl (SRR8362948), and green junglefowl (SRR8362940) respectively at position 3:70297200-70305540 intronic region of *GRIK2* gene is shown. Grey horizontal rectangular boxes represent the reads mapped to the Galgal6 genome assembly, and the white space represents the deleted region in Ceylon junglefowl species. The General Feature Format (GFF/GTF) is loaded at the bottom.

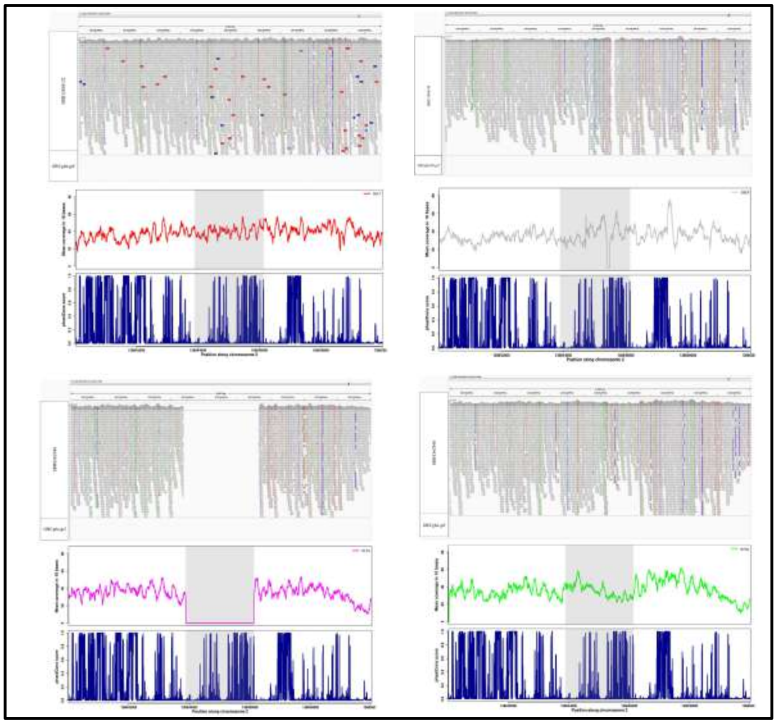

**Fig. S10:** IGV screenshot, mean coverage of 10 bases per window, and phastCons score of the Galgal6 genome assembly of deleted and adjacent regions for four junglefowl species, red junglefowl (SRR12868132), grey junglefowl (SRR12868140), Ceylon junglefowl (SRR8362948), and green junglefowl (SRR8362940) respectively at position 2:138640390-138649640 located in an intergenic region and >30Kb adjacent to *MTSS1* and *SQLE* genes are shown. Grey horizontal rectangular boxes represent the reads mapped to the Galgal6 genome assembly, and the white space represents the deleted region in Ceylon junglefowl species. The General Feature Format (GFF/GTF) is loaded at the bottom.

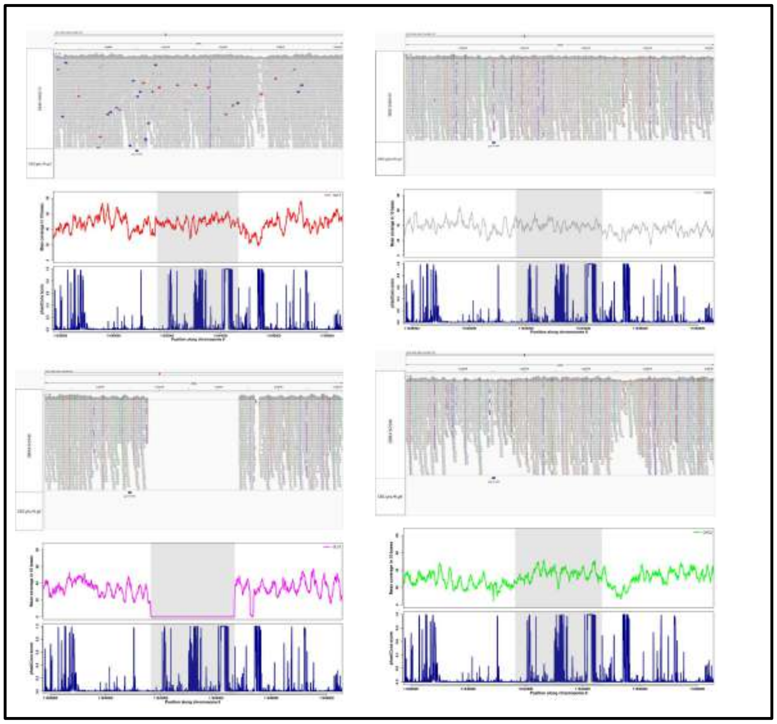

**Fig. S11:** IGV screenshot, mean coverage of 10 bases per window, and phastCons score of the Galgal6 genome assembly of deleted and adjacent regions for four junglefowl species, red junglefowl (SRR12868132), grey junglefowl (SRR12868140), Ceylon junglefowl (SRR8362948), and green junglefowl (SRR8362940) respectively at position 8:11638140-11648170 located in an intergenic region and 49Kbp adjacent to a paralogous copy of *COL11A1* gene are shown. Grey horizontal rectangular boxes represent the reads mapped to the Galgal6 genome assembly, and the white space represents the deleted region in Ceylon junglefowl species. The General Feature Format (GFF/GTF) is loaded at the bottom.

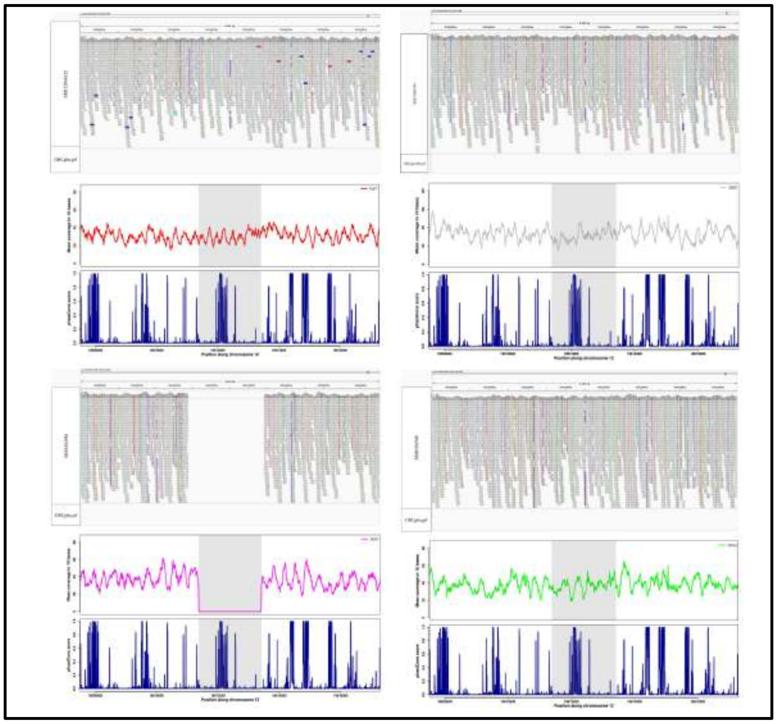

**Fig. S12:** IGV screenshot, mean coverage of 10 bases per window, and phastCons score of the Galgal6 genome assembly of deleted and adjacent regions for four junglefowl species, red junglefowl (SRR12868132), grey junglefowl (SRR12868140), Ceylon junglefowl (SRR8362948), and green junglefowl (SRR8362940) respectively at position 12:19607880-19616920 located in an intergenic region and 16Kb adjacent to *LMCD1* gene are shown. Grey horizontal rectangular boxes represent the reads mapped to the Galgal6 genome assembly, and the white space represents the deleted region in Ceylon junglefowl species. The General Feature Format (GFF/GTF) is loaded at the bottom.

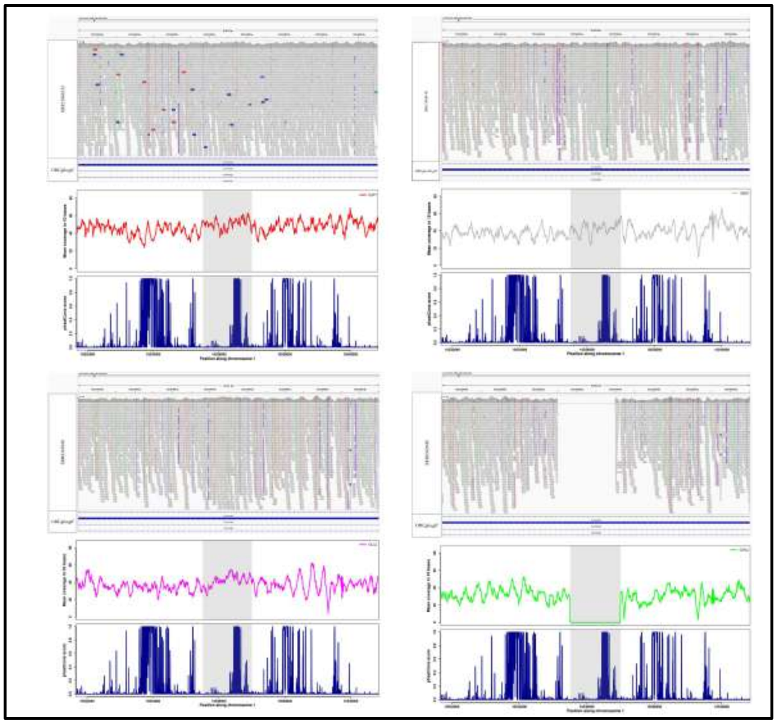

**Fig. S13:** IGV screenshot, mean coverage of 10 bases per window, and phastCons score of the Galgal6 genome assembly of deleted and adjacent regions for four junglefowl species, red junglefowl (SRR12868132), grey junglefowl (SRR12868140), Ceylon junglefowl (SRR8362948), and green junglefowl (SRR8362940) respectively at position 1:10532010-10540500 intronic region of *CACNA2D1* gene is shown. Grey horizontal rectangular boxes represent the reads mapped to the Galgal6 genome assembly, and the white space represents the deleted region in green junglefowl species. The General Feature Format (GFF/GTF) is loaded at the bottom.

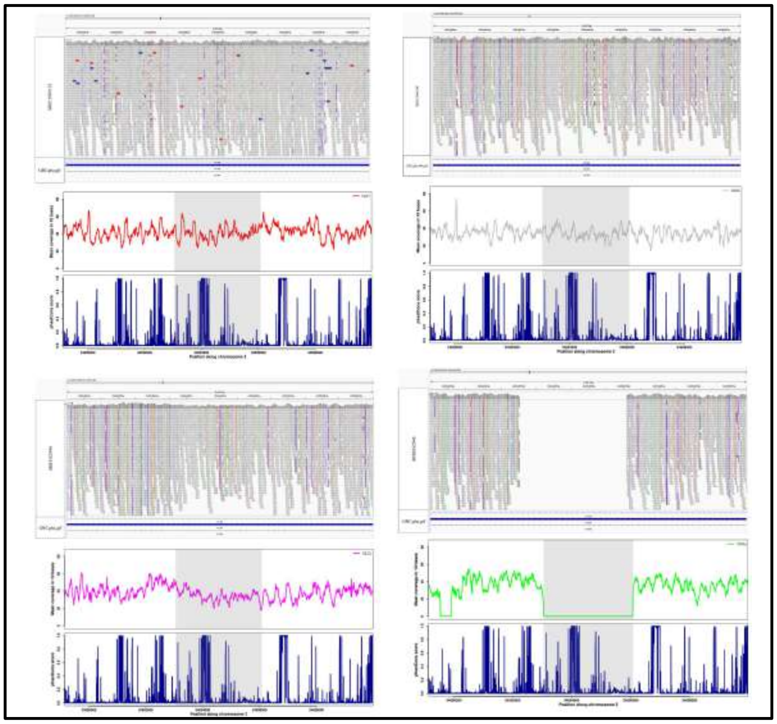

**Fig. S14:** IGV screenshot, mean coverage of 10 bases per window, and phastCons score of the Galgal6 genome assembly of deleted and adjacent regions for four junglefowl species, red junglefowl (SRR12868132), grey junglefowl (SRR12868140), Ceylon junglefowl (SRR8362948), and green junglefowl (SRR8362940) respectively at position 3:34689550-34699580 intronic region of *KIF26B* gene is shown. Grey horizontal rectangular boxes represent the reads mapped to the Galgal6 genome assembly, and the white space represents the deleted region in green junglefowl species. The General Feature Format (GFF/GTF) is loaded at the bottom.

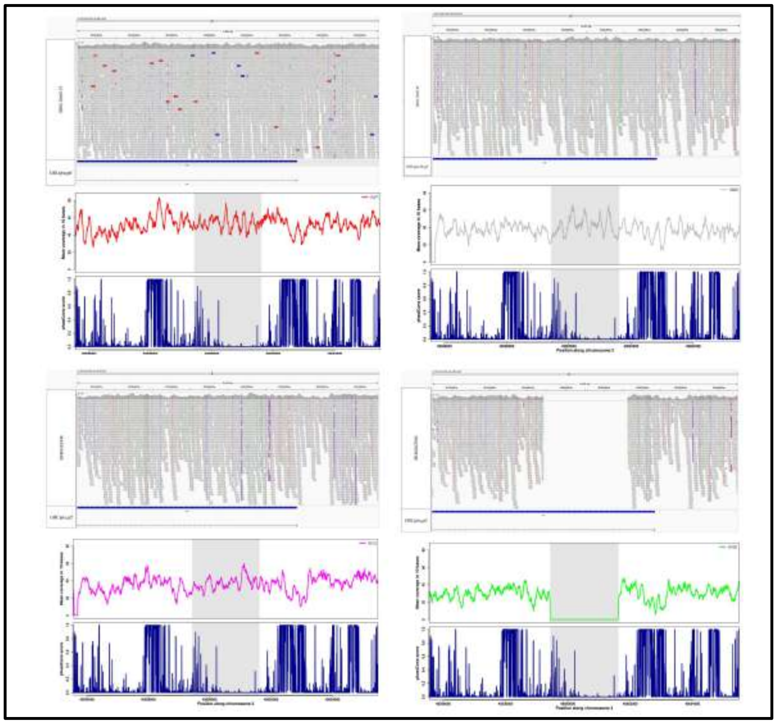

**Fig. S15:** IGV screenshot, mean coverage of 10 bases per window, and phastCons score of the Galgal6 genome assembly of deleted and adjacent regions for four junglefowl species, red junglefowl (SRR12868132), grey junglefowl (SRR12868140), Ceylon junglefowl (SRR8362948), and green junglefowl (SRR8362940) respectively at position 3:49595930-49605120 intronic region of *VIP* gene is shown. Grey horizontal rectangular boxes represent the reads mapped to the Galgal6 genome assembly, and the white space represents the deleted region in green junglefowl species. The General Feature Format (GFF/GTF) is loaded at the bottom.

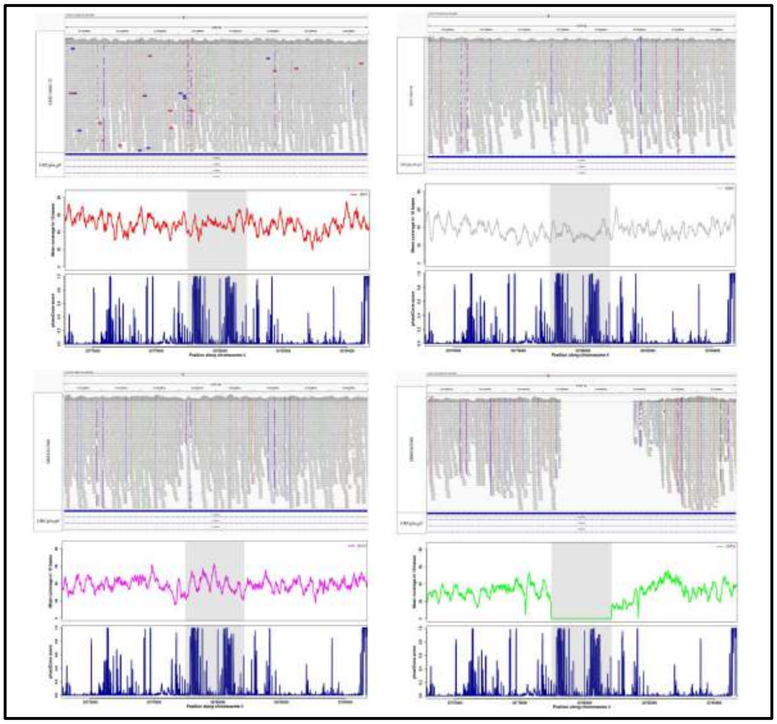

**Fig. S16:** IGV screenshot, mean coverage of 10 bases per window, and phastCons score of the Galgal6 genome assembly of deleted and adjacent regions for four junglefowl species, red junglefowl (SRR12868132), grey junglefowl (SRR12868140), Ceylon junglefowl (SRR8362948), and green junglefowl (SRR8362940) respectively at position 4:35775480-35784340 intronic region of *CCSER1* gene is shown. Grey horizontal rectangular boxes represent the reads mapped to the Galgal6 genome assembly, and the white space represents the deleted region in green junglefowl species. The General Feature Format (GFF/GTF) is loaded at the bottom.

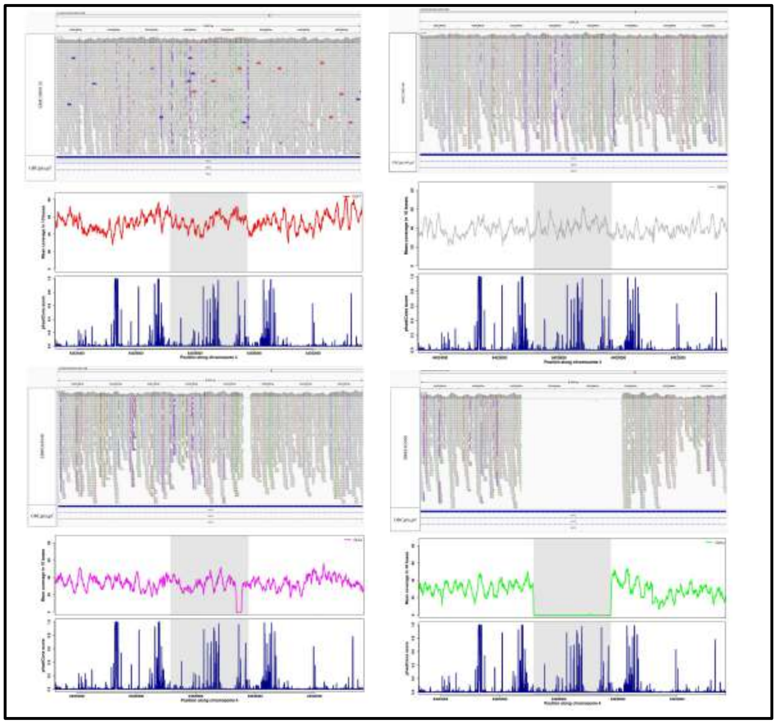

**Fig. S17:** IGV screenshot, mean coverage of 10 bases per window, and phastCons score of the Galgal6 genome assembly of deleted and adjacent regions for four junglefowl species, red junglefowl (SRR12868132), grey junglefowl (SRR12868140), Ceylon junglefowl (SRR8362948), and green junglefowl (SRR8362940) respectively at position 4:64023650-64033270 intronic region of *SGCZ* gene is shown. Grey horizontal rectangular boxes represent the reads mapped to the Galgal6 genome assembly, and the white space represents the deleted region in green junglefowl species. The General Feature Format (GFF/GTF) is loaded at the bottom.

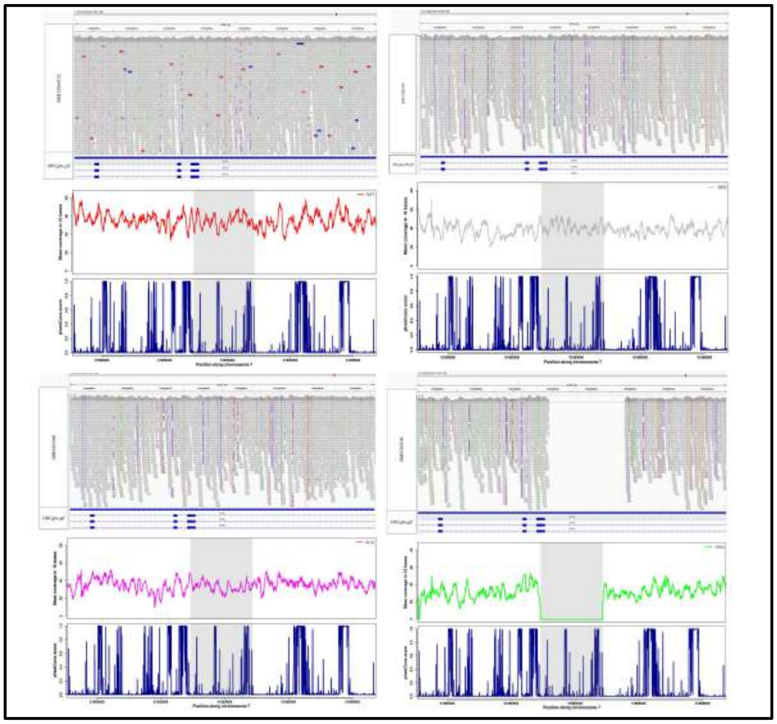

**Fig. S18:** IGV screenshot, mean coverage of 10 bases per window, and phastCons score of the Galgal6 genome assembly of deleted and adjacent regions for four junglefowl species, red junglefowl (SRR12868132), grey junglefowl (SRR12868140), Ceylon junglefowl (SRR8362948), and green junglefowl (SRR8362940) respectively at position 7:31889420-31898370 intronic region of *LRP1B* gene is shown. Grey horizontal rectangular boxes represent the reads mapped to the Galgal6 genome assembly, and the white space represents the deleted region in green junglefowl species. The General Feature Format (GFF/GTF) is loaded at the bottom.

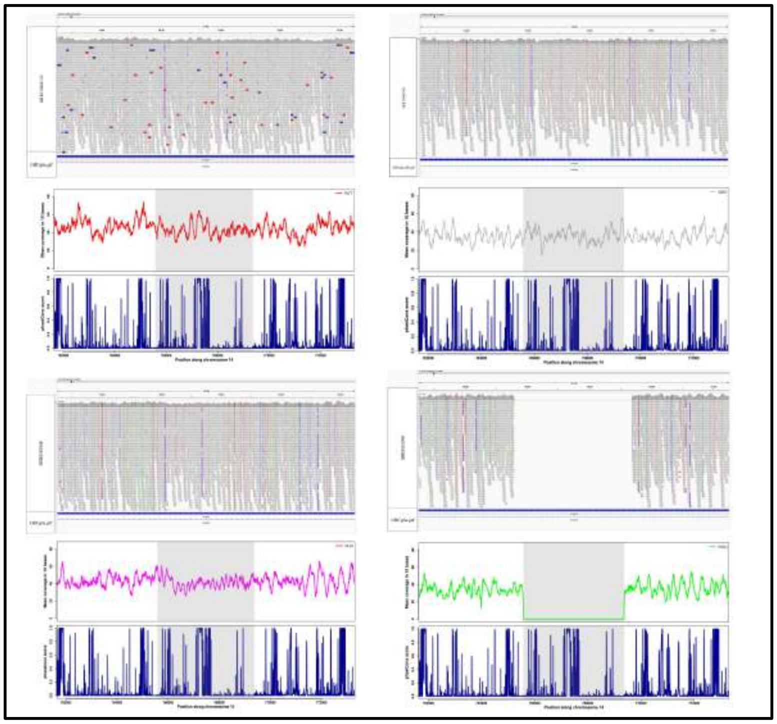

**Fig. S19:** IGV screenshot, mean coverage of 10 bases per window, and phastCons score of the Galgal6 genome assembly of deleted and adjacent regions for four junglefowl species, red junglefowl (SRR12868132), grey junglefowl (SRR12868140), Ceylon junglefowl (SRR8362948), and green junglefowl (SRR8362940) respectively at position 14:762080-772870 intronic region of *SHISA9* gene is shown. Grey horizontal rectangular boxes represent the reads mapped to the Galgal6 genome assembly, and the white space represents the deleted region in green junglefowl species. The General Feature Format (GFF/GTF) is loaded at the bottom.

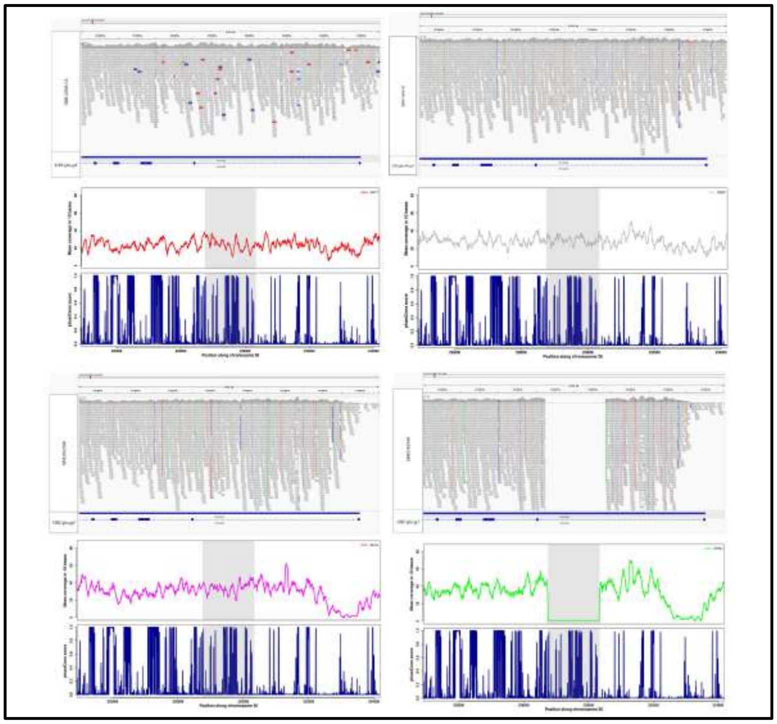

**Fig. S20:** IGV screenshot, mean coverage of 10 bases per window, and phastCons score of the Galgal6 genome assembly of deleted and adjacent regions for four junglefowl species, red junglefowl (SRR12868132), grey junglefowl (SRR12868140), Ceylon junglefowl (SRR8362948), and green junglefowl (SRR8362940) respectively at position 26:225250-233840 intronic region of *TBC1D22B* gene is shown. Grey horizontal rectangular boxes represent the reads mapped to the Galgal6 genome assembly, and the white space represents the deleted region in green junglefowl species. The General Feature Format (GFF/GTF) is loaded at the bottom.

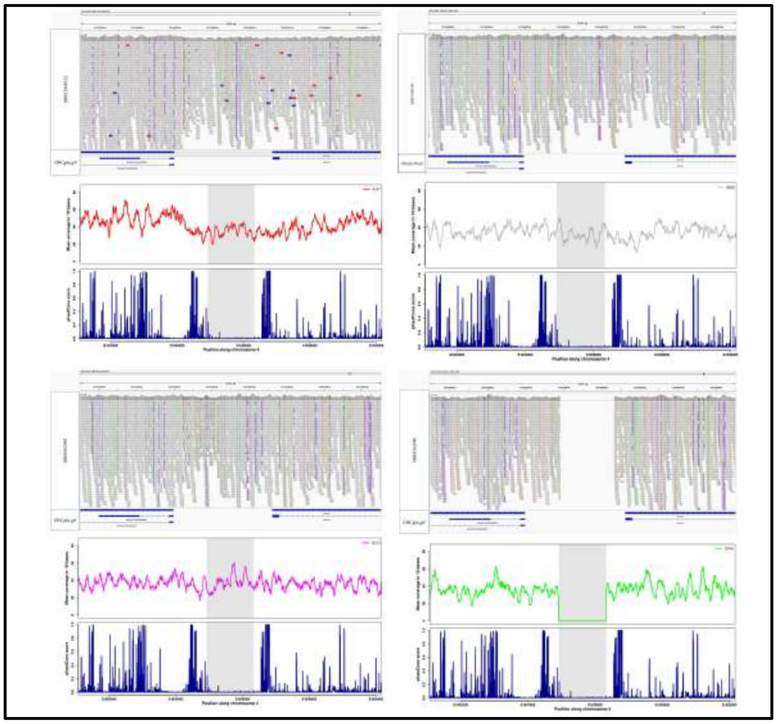

**Fig. S21:** IGV screenshot, mean coverage of 10 bases per window, and phastCons score of the Galgal6 genome assembly of deleted and adjacent regions for four junglefowl species, red junglefowl (SRR12868132), grey junglefowl (SRR12868140), Ceylon junglefowl (SRR8362948), and green junglefowl (SRR8362940) respectively at position 4:81841420-81849830 located in an intergenic region and 600 bp adjacent to *LRPAP1* gene are shown. Grey horizontal rectangular boxes represent the reads mapped to the Galgal6 genome assembly, and the white space represents the deleted region in green junglefowl species. The General Feature Format (GFF/GTF) is loaded at the bottom.

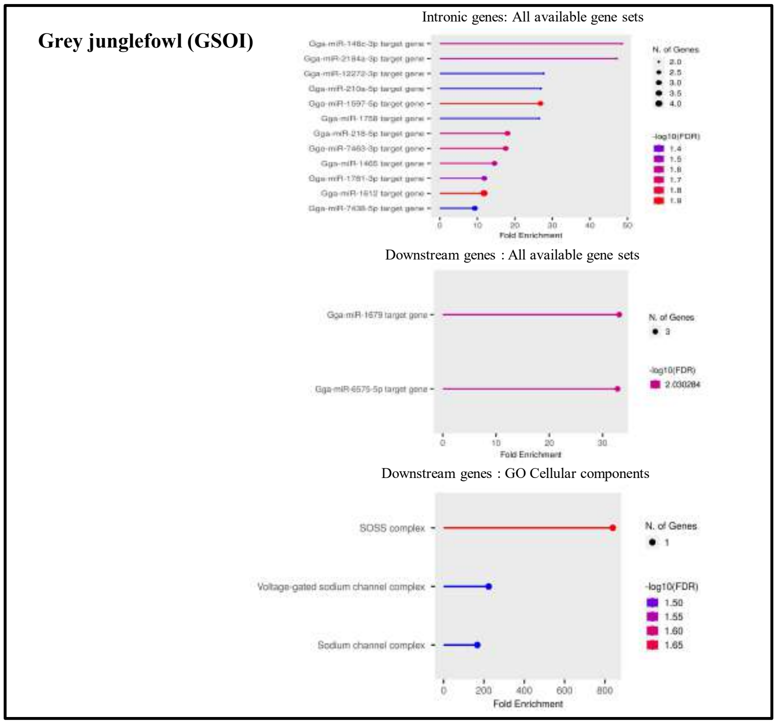


**Fig. S22:** Gene enrichment analysis of the genes with intronic deletions and upstream deletion in grey junglefowl (GSOI) species.

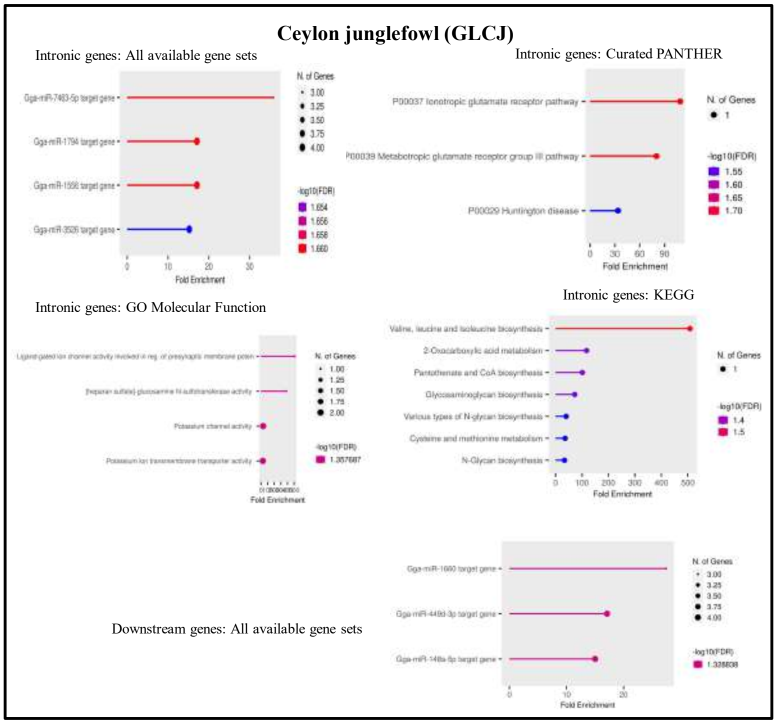


**Fig. S23:** Gene enrichment analysis of the genes with intronic deletions and upstream deletion in Ceylon junglefowl (GLCJ) species.

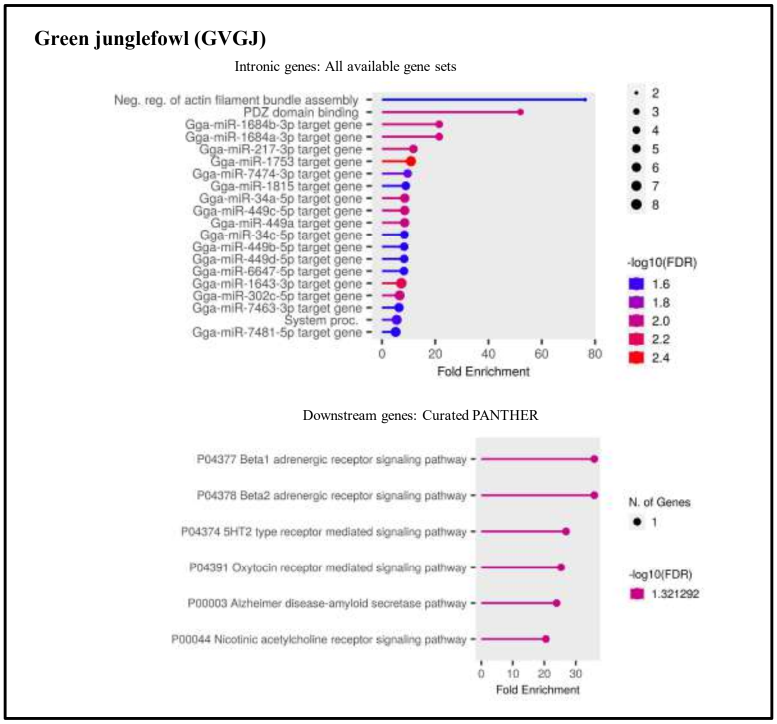


**Fig. S24:** Gene enrichment analysis of the genes with intronic deletions and upstream deletion in green junglefowl species (GVGJ).

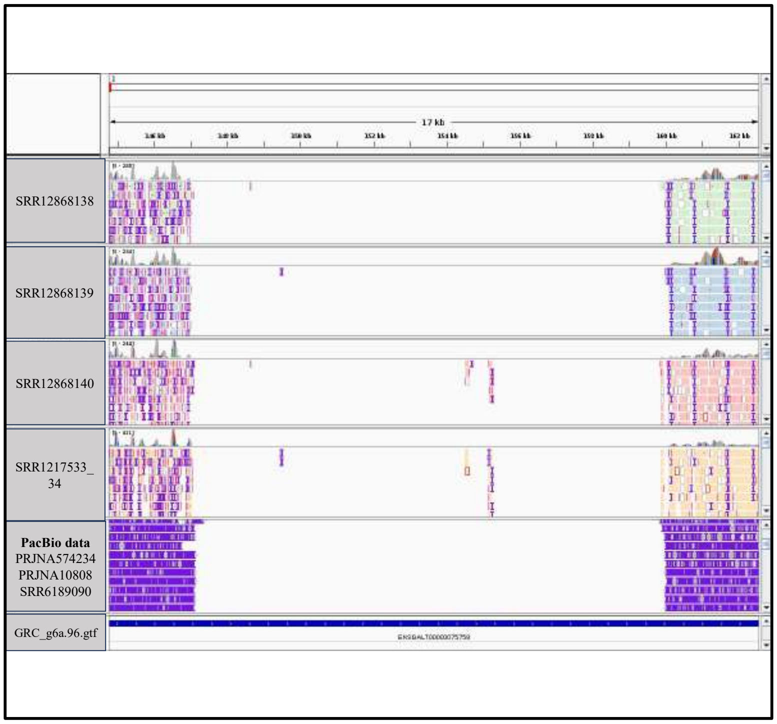


**Fig. S25:** IGV screenshot at position 1:145000-162000 of 3 individuals (SRR12868138, SRR12868139, and SRR12868140) of grey jungle fowl and one individual of red jungle fowl (SRR1217533_34) mapped to Gallus_gallus_GRCg6a genome assembly. Red jungle genome assembly shows a gap in this region. The mapped long-read PacBio data also support this assembly gap at the given position. The General Feature Format (GFF/GTF) is loaded at the bottom.

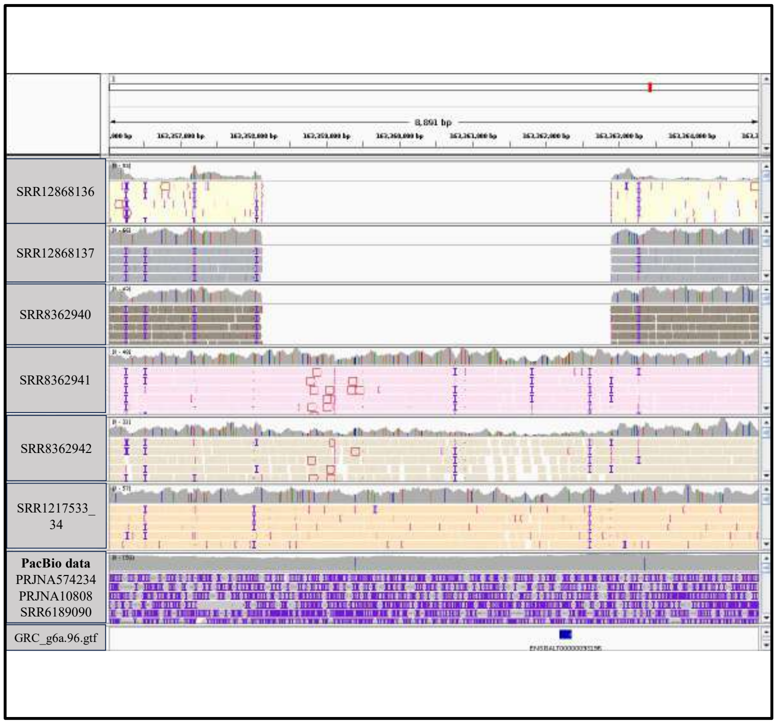


**Fig. S26:** IGV screenshot at position 1:163356000-163364500 of five individuals (SRR12868136, SRR12868137, SRR8362940, SRR8362941, and SRR8362942) of green jungle fowl and one individual of red jungle fowl (SRR1217533_34) mapped to Gallus_gallus_GRCg6a genome assembly. Three individuals of green jungle fowl support a midsize deletion, while the other two individuals are not supporting this midsize deletion. The General Feature Format (GFF/GTF) is loaded at the bottom.

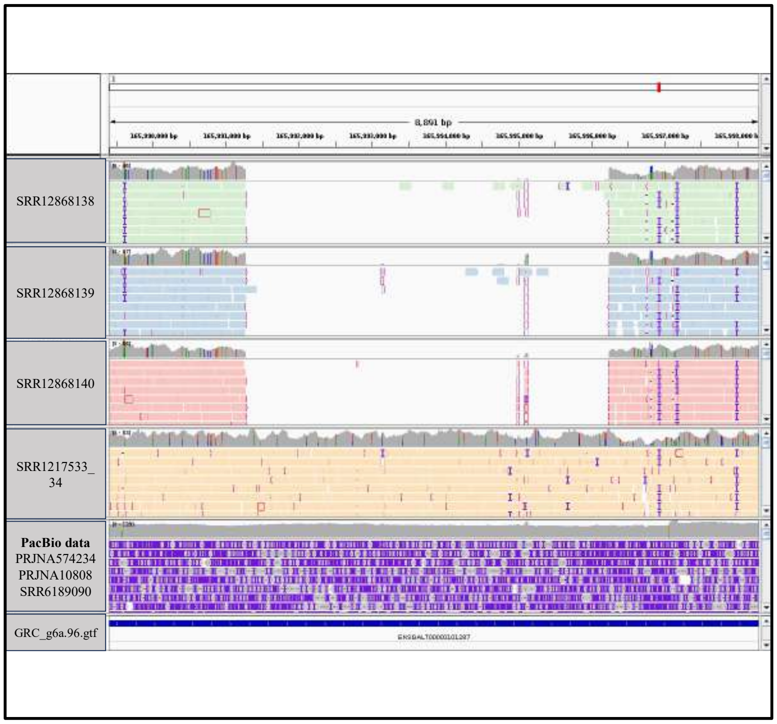


**Fig. S27**: IGV screenshot at position 1:165990000-165998000 for three individuals (SRR12868138, SRR12868139, and SRR12868140) of grey jungle fowl and one individual of red jungle fowl (SRR1217533_34) mapped to Gallus_gallus_GRCg6a genome assembly. Some spurious reads mapped to GRCg6a genome has been found in region of midsize deletion. The General Feature Format (GFF/GTF) is loaded at the bottom.

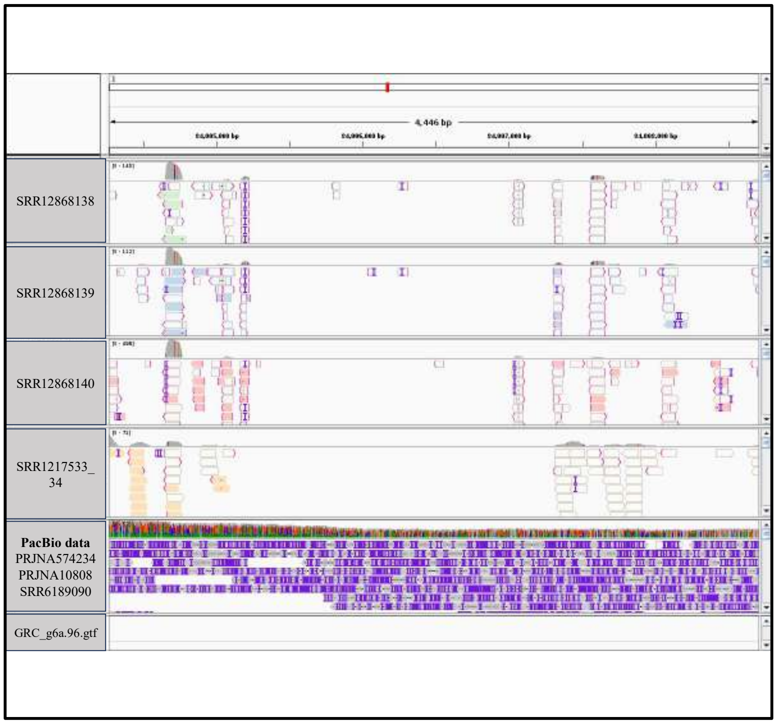


**Fig. S28:** IGV screenshot at position 1:75996000-76000000 for three individuals (SRR12868138, SRR12868139, and SRR12868140) of grey jungle fowl and one individual of red jungle fowl (SRR1217533_34) mapped to Gallus_gallus_GRCg6a genome assembly. Repeats are present in this region in red jungle fowl genome assembly. The General Feature Format (GFF/GTF) is loaded at the bottom.
